# CD8^+^ T-cell memory induced by successive SARS-CoV-2 mRNA vaccinations is characterized by clonal replacement

**DOI:** 10.1101/2022.08.27.504955

**Authors:** Hiroyasu Aoki, Masahiro Kitabatake, Haruka Abe, Peng Xu, Mikiya Tsunoda, Shigeyuki Shichino, Atsushi Hara, Noriko Ouji-Sageshima, Chihiro Motozono, Toshihiro Ito, Kouji Matsushima, Satoshi Ueha

## Abstract

mRNA vaccines against the Spike glycoprotein of severe acute respiratory syndrome type 2 coronavirus (SARS-CoV-2) elicit strong T-cell responses. However, it’s not known whether T cell clonotypes responding to the first vaccination repeatedly expand with booster vaccinations. Here, we temporally tracked the CD8^+^ T-cell repertoire in individuals who received three shots of the BNT162b2 mRNA vaccine. By analyzing the kinetic profile of CD8^+^ T-cell clonotypes responding to the first, second, or third shot, we demonstrated that newly expanded clonotypes elicited by the second shot replaced many of those that responded to the first shot. Although these 2^nd^ responder clonotypes expanded after the third shot, their clonal diversity was skewed, and they were partially replaced by newly elicited the 3^rd^ responders. Furthermore, this replacement of vaccine-responding clonotypes occurred within the same Spike epitope. These results suggest that CD8^+^ T-cell memory induced by repetitive mRNA vaccination is characterized by the emergence of new dominant clones.

## Introduction

mRNA vaccines against the Spike glycoprotein of severe acute respiratory syndrome type 2 coronavirus (SARS-CoV-2), including Pfizer-BioNTech’s BNT162b2 and Moderna’s mRNA-1273, are strikingly effective at reducing coronavirus disease 2019 (COVID-19)^1^. These mRNA vaccines elicit both humoral and cellular immune responses specific for the viral Spike protein^2–4^. The cellular immune response may be more protective to variants of concerns (VOCs) of SARS-CoV-2 than the humoral response, because cellular immunity against the ancestral strain targets relatively conserved regions of the Spike protein^5, 6^ and potentially cross-reacts with the other variants^7, 8^. Studies on the cellular immune responses to SARS-CoV-2 mRNA vaccination have demonstrated that the numbers of Spike-reactive T cells increase in the peripheral blood after the first shot, and are additionally boosted after the second shot^4, 9, 10^. Furthermore, the third mRNA vaccination rapidly re-expands Spike-reactive T cells that had waned after the second shot^11, 12^.

Previous studies demonstrated that secondary, or boost, vaccinations induce the reactivation of memory CD8^+^ T cells established following the initial vaccination^13–16^. However, repetitive antigen stimulation in mice alters the properties of memory T cells, including enhancement of their effector function, and reductions in their proliferation potential and ability to produce IL-2^14, 17^. This suggests that memory CD8^+^ T cells induced by the first vaccination become poorly, or non-responsive during the course of repetitive antigen stimulation. In line with this hypothesis, we previously reported that older memory T cell populations were replaced by newly expanded T cell populations in a serial adoptive transfer model characterized by repetitive antigen exposure^18^.

However, in repetitive SARS-CoV-2 mRNA vaccinations, it is not known whether T cell clonotypes that had responded to the first shot expanded repeatedly after subsequent vaccination, or whether primary responder clonotypes were partially, or fully, replaced by newly expanded clonotypes after each vaccination.

Pathogen-reactive T cells recognize distinct antigenic epitopes via their unique T cell receptor (TCR). To analyze changes in T cell clonal dominance during SARS-CoV-2 mRNA vaccination, we undertook a longitudinal repertoire analysis by TCR sequencing. This enabled us to comprehensively quantify changes in the frequencies of specific T-cell clones (“kinetic profile”) during primary and secondary responses to mRNA vaccination. Specifically, we performed a longitudinal TCR repertoire analysis on peripheral blood T cells at ten time points, before and after the first, second, and third BNT162b2 mRNA vaccination. Furthermore, Spike reactivity, immunological phenotype, and epitope specificity of T cell clonotypes were analyzed by an activation-induced marker assay (AIM assay) using a Spike peptide pool, single-cell analysis of gene expression and TCR sequence, and TCR motif analysis of Spike-specific T-cell clonotypes.

## Results

### Novel clones dominated the T-cell response after the second vaccination

We studied a cohort of 38 healthcare workers at Nara Medical University (Japan) who had received three shots of the Pfizer-BNT162b2 vaccine^19^ (Supplementary Table S1). Peripheral blood mononuclear cell (PBMC) samples were longitudinally collected from the participants at ten points (Fig. 1A). CD4^+^ and CD8^+^ T cells were separated from the PBMC samples, and their TCR repertoire was analyzed (“bulk TCR repertoire analysis”). To analyze the kinetics of T cell clonal responses to mRNA vaccination, we first compared the frequency of clonotypes before and after the vaccination (P1-P2, P2-P3, and P6-P8). We then identified clonotypes that significantly expanded after each vaccination (i.e., vaccine-responding clonotype) (Figure 1B)^20^. For both CD4^+^ and CD8^+^ T cells, the total frequency and diversity of clonotypes that expanded after the second shot was greater than that of responding clonotypes to the first and third shots (Figure 1C and D). This result demonstrated that the T cell clonal response to mRNA vaccination peaked after the second shot.

**Fig. 1.**
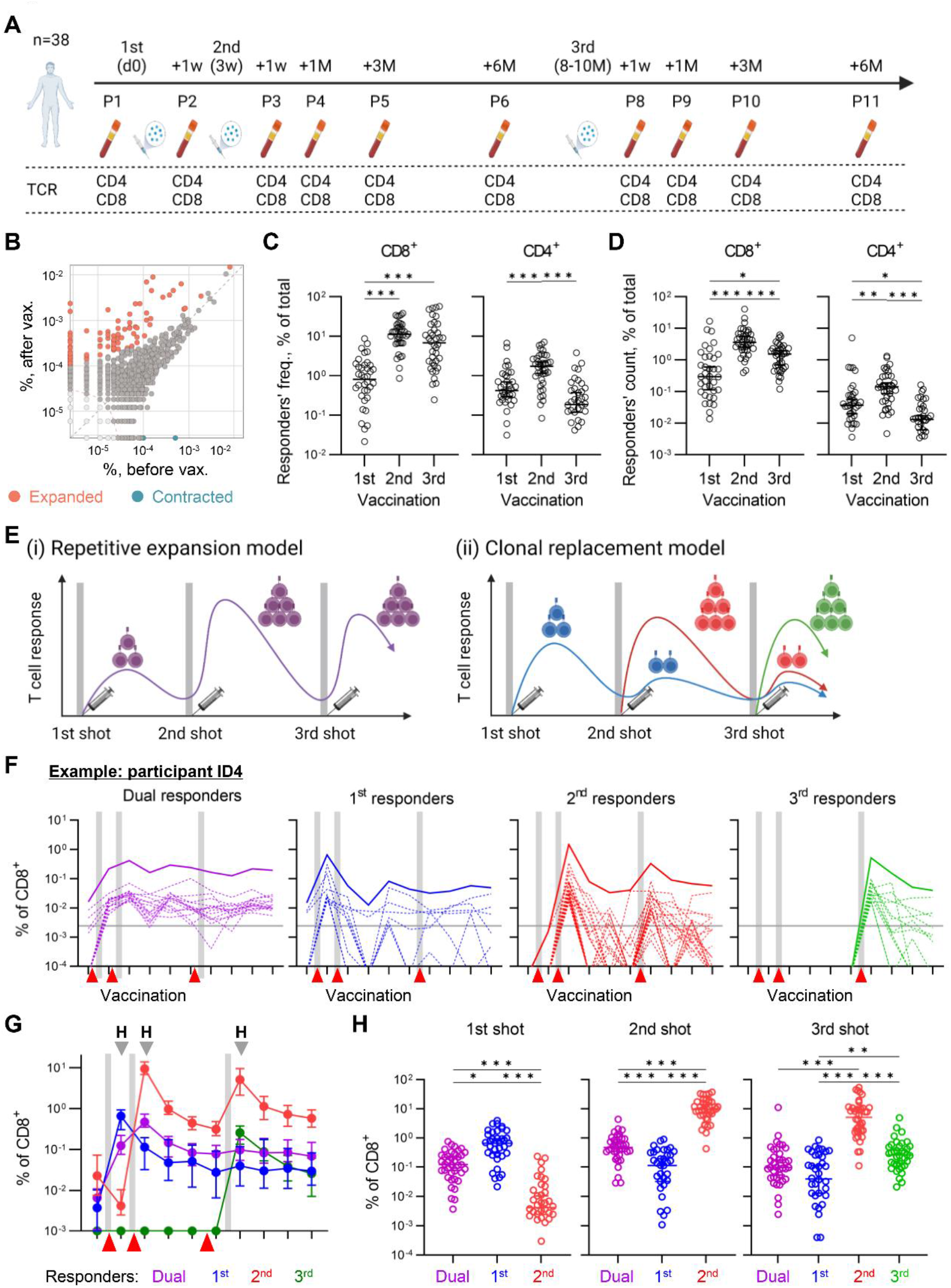
Identifying responding clonotypes after vaccination. (A) Schematic of longitudinal sample collection before and after BNT162b2 vaccination. Time points for Bulk TCR-seq (TCR). (B) Example for identifying responding clonotypes using the beta-binomial model^20^. The scatter plot represents clonotype frequencies in samples collected before and after vaccination. Significantly expanded (responding) and contracted clonotypes were indicated as orange and cyan. (C) The total frequency of CD8^+^ (left) and CD4^+^ (right) responding clonotypes after each vaccination. (D) Diversity of CD8^+^ (left) and CD4^+^ (right) responding clonotypes after each vaccination. The count of responding clonotypes normalized by the total number of clones for each participant was plotted. (E) Models for T-cell responses following mRNA vaccinations. (left) Repetitive expansion model: the clonotypes responded to the first shot and expanded repeatedly after the second and third shots. (right) Clonal replacement model: primary responder clonotypes were turned over after each vaccination shot. (F) Classifying CD8^+^ T-cell clonotypes into 1^st^, 2^nd^, dual, and 3^rd^ responders. The frequency of each clonotype was plotted longitudinally for each type of reaction (dashed lines). The longitudinal changes of their total frequency were also plotted (thick line). The red arrows represent the timing of vaccination. The grey horizontal line represents the threshold for a clonotype to be regarded as “Increased.” The data from one participant (ID4) is shown. (G) Kinetics of CD8^+^ 1^st^, 2^nd^, dual, and 3^rd^ responder clone frequencies. The median with 95% CI was plotted. (H) The comparison of the total frequency of CD8^+^ 1^st^, 2^nd^, dual, and 3^rd^ responder clonotypes after each shot. The total frequency for 3^rd^ responders was plotted only after the third shot (right). (C, D, H) Median was plotted. Friedman test with Dunn’s multiple comparison. n = 38 except for P11 (n=37). ***P<0.001; **P<0.01; *P<0.05. (A) and (E) were created using BioRender.com.

Using the longitudinal bulk TCR-seq data, we analyzed which of the following two models explained the T-cell responses following mRNA vaccinations: (i) repetitive expansion model, in which the clonotypes responded to the first shot expanded repeatedly after the second and third shot, or (ii) clonal replacement model, in which primary responder clonotypes were partially or fully replaced by newly expanded ones after each shot (Figure 1E). For this purpose, the responding clonotypes were classified into four patterns based on their response to each shot (Figure 1F and Figure S1A): 1^st^ responders (expanded after the first shot but did not after the second shot), 2^nd^ responders (did not expand after the first shot but expanded after the second shot), dual responders (expanded or increased after both the first and the second shot), and 3^rd^ responders (were not detected before the third shot and responded to the third shot) (Materials and Methods). Then, the clonal response of each subject was summarized by calculating the total frequency of the clonotypes in each response pattern (Figure 1F, bold line). We longitudinally tracked the total frequency of each response pattern (Figure 1G). After the first shot, the total frequency of the 1^st^ and dual responders was significantly higher than that of the 2^nd^ responders. On the other hand, after the second shot, the total frequency of 2^nd^ responders exceeded the 1^st^ and dual responders (Figure 1H). The fact that the 2^nd^ responders, not the dual responders, were dominant after the second shot suggests that, after the second shot, the majority of the responding clonotypes were new clonotypes. The total frequency of responders after the third shot was also analyzed (Figure 1H). At this point, although the expansion of 3^rd^ responders was observed, the 2^nd^ responders were still dominant, and their total frequency was more than twenty times that of 3^rd^ responders. This result demonstrated that most of the responding clonotypes were maintained between the second and third shots, unlike the predominant replacement observed between the first and second shots. A similar replacement of responding clones was also observed in CD4^+^ T cells (Figure S1A and B).

### 2^nd^ responders were more dominant in CD8+ Spike-reactive clonotypes than dual responders

Next, we analyzed whether the predominant replacement of responding clonotypes between the first and the second vaccination was also observed among Spike-reactive clones (Figure 2A). To detect Spike-reactive clonotypes, an activation-induced marker (AIM) assay was performed^9, 21–23^. PBMCs collected after the vaccinations (P2, P3, and P8) were stimulated with peptide pools of the SARS-CoV-2 Spike protein. Then, CD137^+^ or CD107a^+^ non-naïve CD8^+^ T cells and CD40L^+^ CD69^+^ non-naive CD4^+^ T cells were sorted as AIM^+^ Spike-reactive cells (Figure 2B and Figure S2A). The proportion of AIM^+^ cells among CD8^+^ and CD4^+^ T cells quantified by flow cytometry peaked after the second shot, consistent with the numbers and extent of expansion of responding clonotypes in the CD8^+^ and CD4^+^ T-cell repertoire (Figure S2B, Figure 1C, and 1D). We then performed TCR-seq on AIM^+^ and AIM^-^ non-naive T cells from the samples after the second shot (P3) and identified AIM^+^ CD8^+^ and CD4^+^ T cell clonotypes (“AIM TCR-seq,” Figure 1A, 2C, and D). The total frequency of AIM^+^ clonotypes was significantly increased, not only after the second shot, but also after the third shot (Figure 2E and Figure S2C). This observation confirms that CD8^+^ and CD4^+^ AIM^+^ clonotypes include Spike-reactive clonotypes responded to the vaccination.

**Fig. 2.**
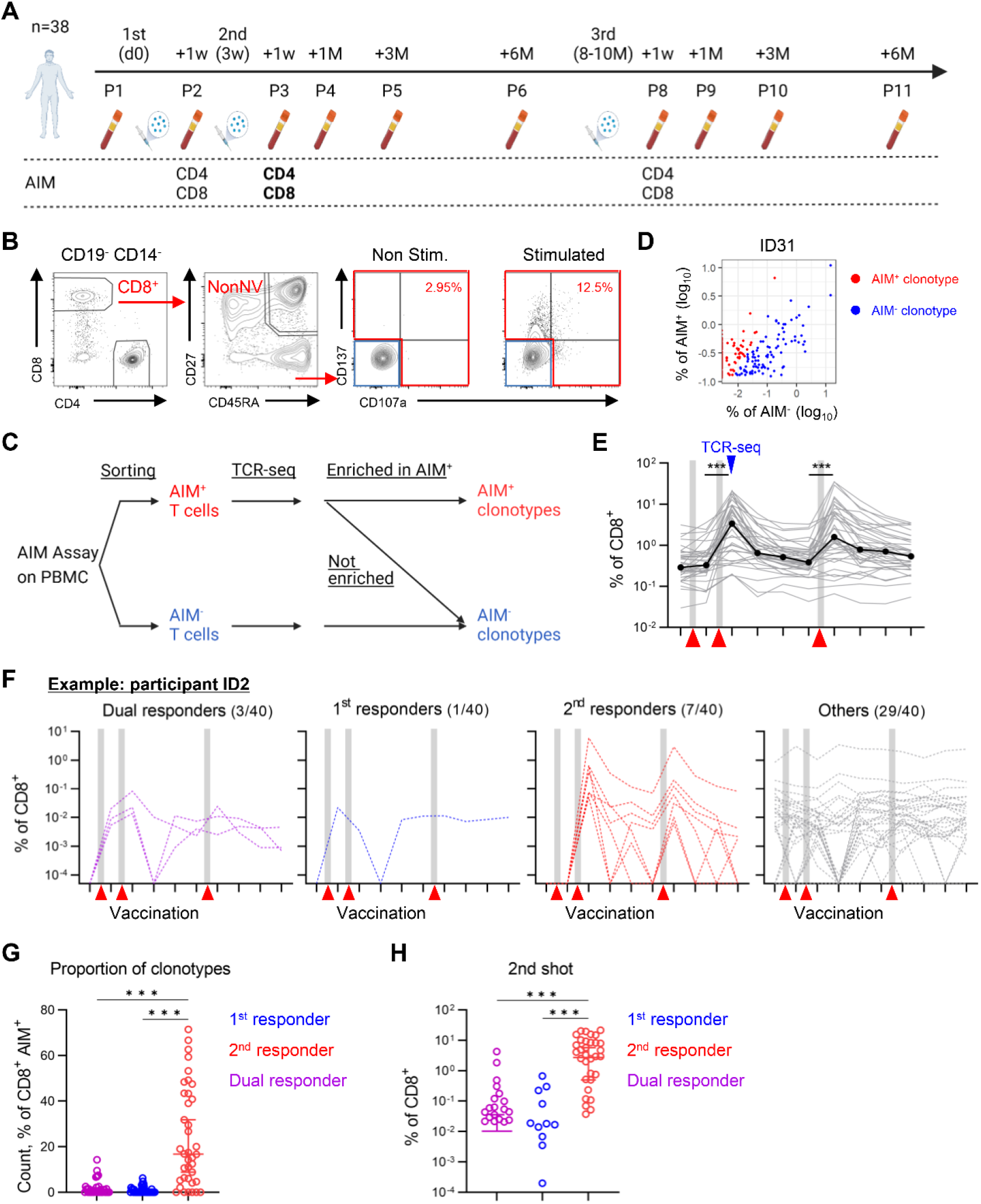
Response pattern of CD8^+^ Spike-reactive clonotypes. (A) Schematic of time points of AIM-assay. AIM TCR-seq was performed only on P3 (bold). (B) Gating strategy for sorting AIM^+^ and AIM^-^ non-naive CD8^+^ T cells. The proportion of AIM^+^ cells among non-naive CD8^+^ T cells was also shown. (C) Flow chart for identifying AIM^+^ clonotypes. AIM^+^ and AIM^-^ T cells were sorted using the gating as (B), then TCR-seq was performed on these T cell subsets. AIM^+^ clonotypes were identified as clonotypes that were enriched in the AIM^+^ T-cell repertoire (Materials and Methods). (D) Scatter plot of overlapping clones between AIM^+^ and AIM^-^ repertoire. Individual dots represent clonotypes with their frequency in AIM^+^ and AIM^-^ repertoire. Red dots represent AIM^+^ clonotypes. (E) Kinetics of the total frequency of CD8^+^ AIM^+^ clonotypes of each participant (thin line, grey). The median was also plotted (thick line, black). (F) Longitudinal tracking of individual CD8^+^ AIM^+^ clonotypes in one participant (ID2). Clonotypes were plotted by response patterns (1^st^ responders: blue, 2^nd^ responders: red, dual responders: purple, and other clonotypes: gray). Numbers indicate the count of the AIM^+^ responder clonotypes and that of all AIM^+^ ones. (G) The proportion of 1^st^, 2^nd^, and dual responders in CD8^+^ AIM^+^ clonotypes. The count of AIM^+^ responder clonotypes divided by that of all AIM^+^ clonotypes was compared. (H) The total frequency of CD8^+^ AIM^+^ 1^st^, 2^nd^, and dual responders after the second vaccination. (E, G, and H) Friedman test with Dunn’s multiple comparisons. n = 38 except for P11 (n=37). ***P<0.001; **P<0.01. (A and C) were created using BioRender.com.

We next classified AIM^+^ clonotypes into the response patterns described above (Figure 2F). In addition, we calculated the proportions of 1^st^, 2^nd^, and dual responders among CD8^+^ AIM^+^ clonotypes (i.e., the number of AIM^+^ 1^st^, 2^nd^, or dual responder clonotypes divided by the total number of AIM^+^ clonotypes). More than half of the CD8^+^ AIM^+^ clonotypes were not classified as responders, suggesting that some non-Spike-reactive bystander clonotypes contaminated the AIM^+^ clonotypes. At the same time, the proportion of the CD8^+^ 2^nd^ responders was significantly higher than that of AIM^+^ dual responders (Figure 2G). Moreover, the total frequency of AIM^+^ 2^nd^ responders of CD8^+^ T cells was also considerably greater than that of AIM^+^ dual responders after the second shot (Figure 2H). These results indicate that the predominant replacement of responding clonotypes between the first and the second shot also occurred among Spike-reactive T cells. Similar results were observed for CD4^+^ AIM^+^ T cell clonotypes (Figure S2D and E).

### 2^nd^ responder clonotypes exhibit effector memory phenotype after the second and third vaccination

Our data demonstrate that 2^nd^ responder clonotypes predominate in T cell responses after the second and the third vaccination. Therefore, we next analyzed the phenotype of 2^nd^ responders after the second and the third vaccination. Targeted single-cell RNA-sequencing (scRNA-seq), single-cell TCR sequencing (scTCR-seq), and protein expression analysis using oligo-conjugated antibodies (AbSeq) were performed on blood CD8^+^ T cells of seven participants collected after the second or third vaccination (P3 and P8) (Figure 3A, S3A, and Supplementary Table S2). After removing γδ T cells and mucosal-associated invariant T cells (MAITs), we categorized conventional non-naïve CD8^+^ T cells into six clusters using integrated mRNA and surface protein expression data (Figure 3B and Figure S3A). We then evaluated the characteristics of these clusters based on the mRNA and protein expression levels of marker genes (Figure 3C, 3D, S3B, and S3C) and signature scores for previously reported blood CD8^+^ T cell clusters^24^ and CD8^+^ T cell characteristics (BD Immune response panel, Hs) (Figure S3D and S3E). Cluster 3 cells expressed *TCF7*, CCR7, and CD127, and exhibited a higher score for stem-like and progenitor-exhausted T cells, which were considered to be central memory T cells (T_CM_). In addition, we regarded Cluster 2 cells as effector-memory T cells (T_EM_), because they expressed *CCR5*, *CD38*, and HLA-DR, and exhibited a higher score for EM-like T cells. Moreover, this cluster included *TOP2A^+^*proliferating T cells. Cells in the remaining four clusters (#0, 1, 4, and 5) were CD45RA^high^, *GZMB*^high^, and CCR7^low^, and exhibited higher effector function scores, matching the characteristics of effector memory T cells expressing CD45RA (T_EMRA_)^25, 26^. Among them, cluster 0 cells were terminal effector (T_TE_), based on their expression of CD56 and *GNLY*, and clusters 4 and 5 cells were named OX40 (CD134) ^+^ and Tim3^+^ by their surface marker expression, respectively. Finally, we named cluster 1 cells transitory (Trans.) T cells because this cluster exhibited an intermediate phenotype between T_EM_ and T_TE_ clusters. The clonality of these CD8^+^ T-cell clusters was calculated from the size distribution of clonotypes determined by scTCRseq. These data indicated that clusters 0 and 1 were primarily comprised of oligoclonal T cells, while cluster 3 was primarily comprised of polyclonal T cells (Figure S3F). This tendency in clonality was consistent after the second and third vaccinations.

**Fig. 3.**
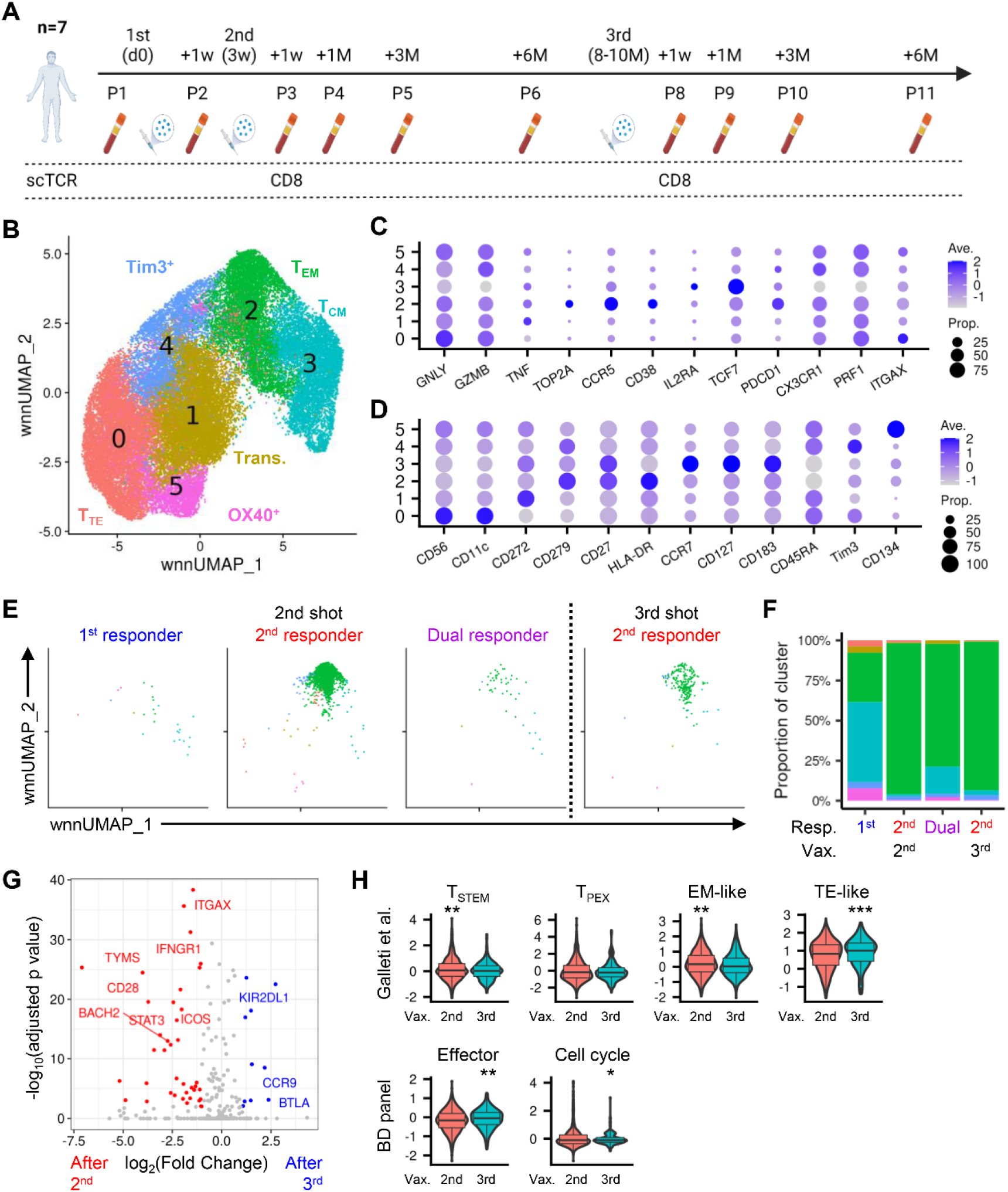
Single-cell analysis on non-naive CD8^+^ T cells after vaccination. (A) Schematic of time points of single-cell analysis. (B) Weighted-nearest neighbor graph-based uniform manifold approximation and projection (wnnUMAP) plot of the scRNAseq/AbSeq data of non-naive CD8^+^ T cells. (C and D) Dot plot showing mRNA (C) and protein (D) expression of marker genes in the T-cell clusters. The size of the circle shows the percentage of cells in a cluster expressing the gene: color scale shows the gene expression level. (E) wnnUMAP plot for T cells in 1^st^, 2^nd^, and dual responder clones at TP3 or TP8. (F) Bar graph representing the cluster distribution of T cells in 1^st^, 2^nd^, and dual responder clonotypes. (G) Volcano plots of differential expression test of T cells from TP3 versus TP8 2^nd^ responder clonotypes. Genes expressed more than two-fold with statistical significance (p < 0.01) after the second or third shot are plotted in red or blue, respectively. (H) Violin plot showing gene signature scores of T cells in 2^nd^ responder clonotypes at TP3 or TP8. Signature genes are referred from the marker genes of T-cell clusters described in Galleti et al.^24^ and the datasheet of BD Rhapsody™ Immune Response Panel Hs. (H) Student’s t-test was performed, and their q-values were estimated. T_TE_: terminal effector T cell; Trans: transitional; T_EM_: effector memory T cell; T_CM_: central memory T cell; T_STEM_: stem-like T cell; T_PEX_: progenitor exhausted-like T cell. ***q<0.001; **q<0.01; *q<0.05. (A) was created using BioRender.com.

To analyze the phenotype of 2^nd^ responder clonotypes, the 1^st^, 2^nd^, and dual responder T cells were plotted in a dimensionality reduction plot (Figure 3E). This analysis revealed that over 90% of T cells in 2^nd^ responder clones exhibited a T_EM_ phenotype after the second shot. On the other hand, the proportion of T_EM_ cells was lower in 1^st^ and dual responders (30. 8% and 76.6 %, respectively) (Figure 3F). Furthermore, the proportion of T_EM_ in the 2^nd^ responders was nearly unchanged after the third shot (94.2% after the second shot, and 92.7 % after the third shot) (Figure 3F). These results demonstrate that 2^nd^ responders are mainly comprised of T_EM_ cells after the second and third vaccination. We also identified differentially expressed genes in 2^nd^ responder T_EM_ cells between the second and third shots to examine their shift of gene expression profile (Figure 3G).

Genes whose expression was significantly decreased after the third shot included genes associated with the cell cycle (*TYMS* and *MCM4*), transcription factors associated with memory formation (*BACH2* and *STAT3*)^27, 28^, and costimulatory receptors (*CD28*, *ICOS*, and *DPP4* (coding CD24))^29, 30^. Moreover, T_EM_ cells in the 2^nd^ responder pool exhibited higher scores for the TE-like T cells and effector function after the third shot (Figure 3H). These results suggest that, although still exhibiting T_EM_ phenotype, 2^nd^ responders are more differentiated towards a T_EMRA_-like phenotype after the third shot.

### Skewed re-expansion of 2^nd^ responder clonotypes after the third shot

Next, we analyzed the changes in the repertoire of 2^nd^ responders after the second and the third shot in more detail. The total frequency and fold expansion of 2^nd^ responders after the third shot was significantly decreased compared to that after the second shot (Figure 4A and S4A), suggesting that the expansion of 2^nd^ responders was diminished after the third shot. Notably, the count of expanded clonotypes after the third shot within 2^nd^ responders (“re-expanded 2^nd^ responders”) was significantly lower than the total count of 2^nd^ responders (Figure 4B): the proportion of re-expanded 2^nd^ responders was only about 30% (Figure S4B). These re-expanded 2^nd^ responder clonotypes were present at a higher frequency after the second shot than 2^nd^ responders that did not expand (“Unchanged”) or were not detected after the third shot (Figure S4C). In line with the reduction of expanding clonotypes among 2^nd^ responders, the relative distribution of 2^nd^ responder clonotypes was also changed after the third shot: their clonality was increased while the evenness of their distribution was decreased (Figure 4D). On the other hand, the appearance of the 3^rd^ responders (about 5% of 2^nd^ responder clonotypes, Figure S4B), partially rescued the decreased diversity of the 2^nd^ responders. These changes in CD8^+^ 2^nd^ responders were also observed in CD4^+^ T cell repertoire (Figure S4D-H). Based on these results, we modeled the response of CD8^+^ T-cell clonotypes to mRNA vaccination (Figure 4E). After the second shot, novel responding clonotypes (2^nd^ responders) largely replaced the clonotypes that expanded after the first shot (1^st^ responders) and dominated the responses. After the third shot, a subset of 2^nd^ responders expanded again resulting in a narrowing of the repertoire (i.e. a reduction in the number of re-expanded clonotypes). At the same time, 3^rd^ responders appeared after the third shot and partially compensated for the decrease in 2^nd^ responders’ diversity.

**Fig. 4.**
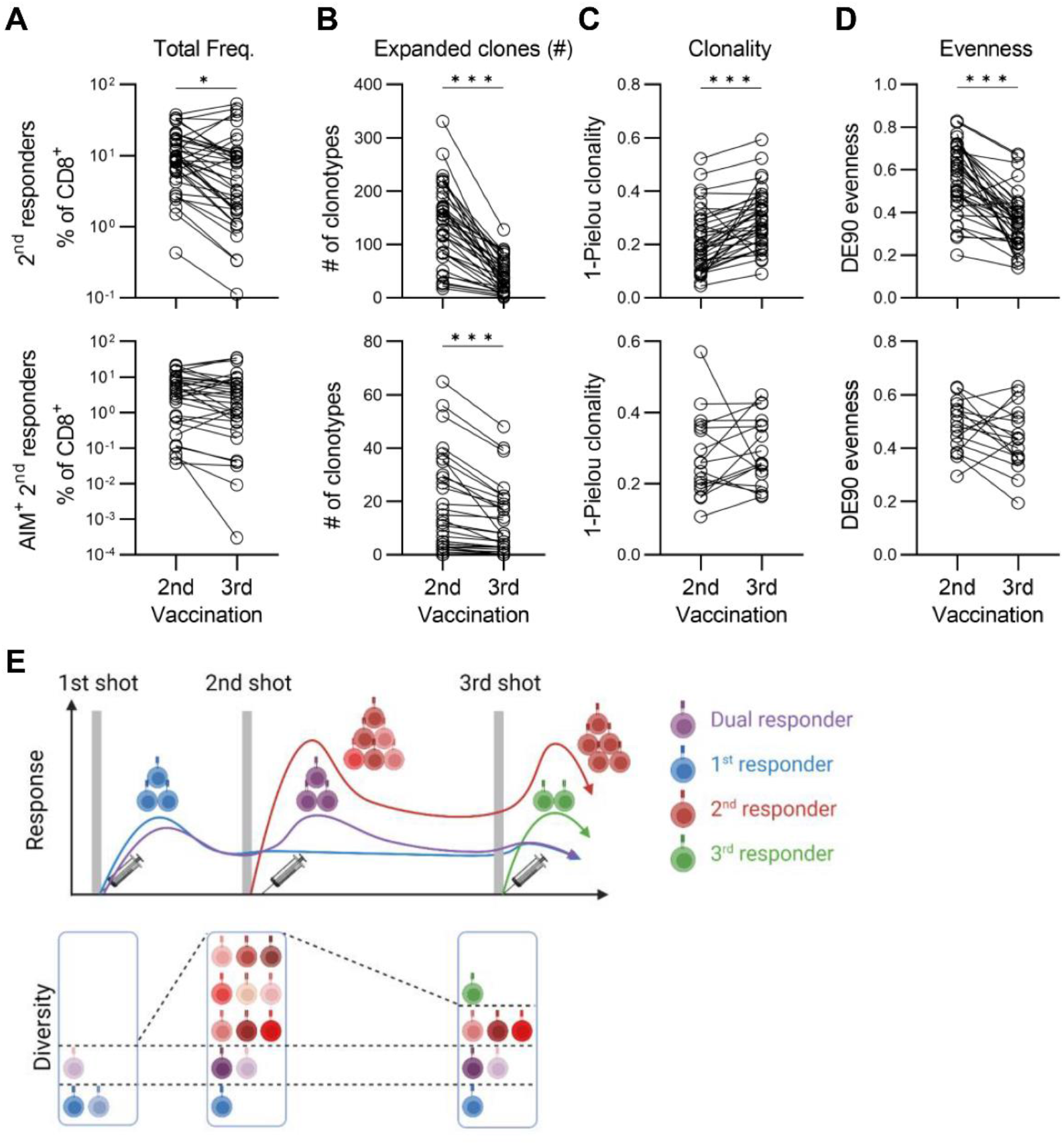
Re-expansion of CD8^+^ 2^nd^ responder clonotypes after the third shot. (A) The total frequency of all (top) or AIM^+^ (bottom) 2^nd^ responders after the second or the third shot. (B) Count of clonotypes that expanded after the second or the third shot within all (top) or AIM^+^ (bottom) 2^nd^ responders. (C) Clonality of all (top) or AIM^+^ (bottom) 2^nd^ responders after the second or the third shot. (D) Evenness of all (top) or AIM^+^ (bottom) 2^nd^ responders after the second or the third shot. (E) Schematic overview for the response of CD8^+^ T-cell clonotypes following mRNA vaccination. After the second shot, 2^nd^ responders turned over 1^st^ responders and became dominant in vaccine-responding T cells. After the third shot, 2^nd^ responders expanded again, while their diversity was reduced. 3^rd^ responders also appeared after the third shot and partially replenished the vaccine-induced T-cell responses. (A-D) Wilcoxon matched-pairs signed rank test. (C and D) Clonality and evenness are calculated only when more than ten clonotypes exist. n = 38 except for the bottom of (C) and (D) (n=17). ***P<0.001; *P<0.05. (E) was created using BioRender.com.

### Induction of CD8^+^ 2^nd^ responders was associated with anti-RBD antibody titers induced by the vaccination

Next, we analyzed the association between the T cell repertoire and other variables, including the participants’ background, anti-SARS-CoV-2 Spike receptor-binding domain (RBD) IgG antibody titer, and strength of the adverse events (Figure 5A). Before doing so, we identified mutually correlated repertoire parameters (e.g., the total frequency of CD8^+^ dual responder at P2, P3, and P8) and merged them into repertoire meta-parameters (e.g., CD8^+^ Dual Responders: DualResp) (Figure 5B and S5A, Materials and Methods). The correlation heatmap between CD8^+^ repertoire meta-parameters and variables related to vaccine response revealed that the total frequency of 2^nd^ responders after the second and the third shot (“2^nd^Resp_P3” and “2^nd^Resp_P8”) was positively correlated with the anti-RBD antibody titers after the second and the third shot, respectively (Figure 5C). Notably, 2^nd^Resp_P3 was significantly correlated with the antibody titer to RBD of the Omicron variant (Figure 5D). These results imply that some participants respond more readily to mRNA vaccination. At the same time, the total frequency of 1^st^ responders (“1^st^Resp”) was positively associated with the severity of adverse events after the vaccination (Figure 5C). As for CD4^+^ T cells, the positive correlation between the repertoire meta-parameters and the severity of adverse events was also shown (Figure S5B). These observations are consistent with the study reporting the association between the T-cell response and the severity of adverse reactions following mRNA-1273 vaccination^31^. Finally, we observed a positive correlation between the related meta-parameters between CD8^+^ and CD4^+^ T-cell repertoire; CD8^+^ 3^rd^Resp and CD4^+^ 3^rd^Resp, CD8^+^ ExpDiv_P8 and CD4^+^ 2^nd^Resp_P8, CD8^+^ AIM and CD4^+^ AIM or 2^nd^Resp_AIM, and so on (Figure S5C). This result implies that the clonal responses of CD4^+^ and CD8^+^ T cells to the mRNA vaccine are partly coordinated. Notably, we observed no significant association between CD4^+^ and CD8^+^ repertoire meta-parameters and sex, history of SARS-CoV-2 infection, or the presence of specific HLA genotypes.

**Fig. 5.**
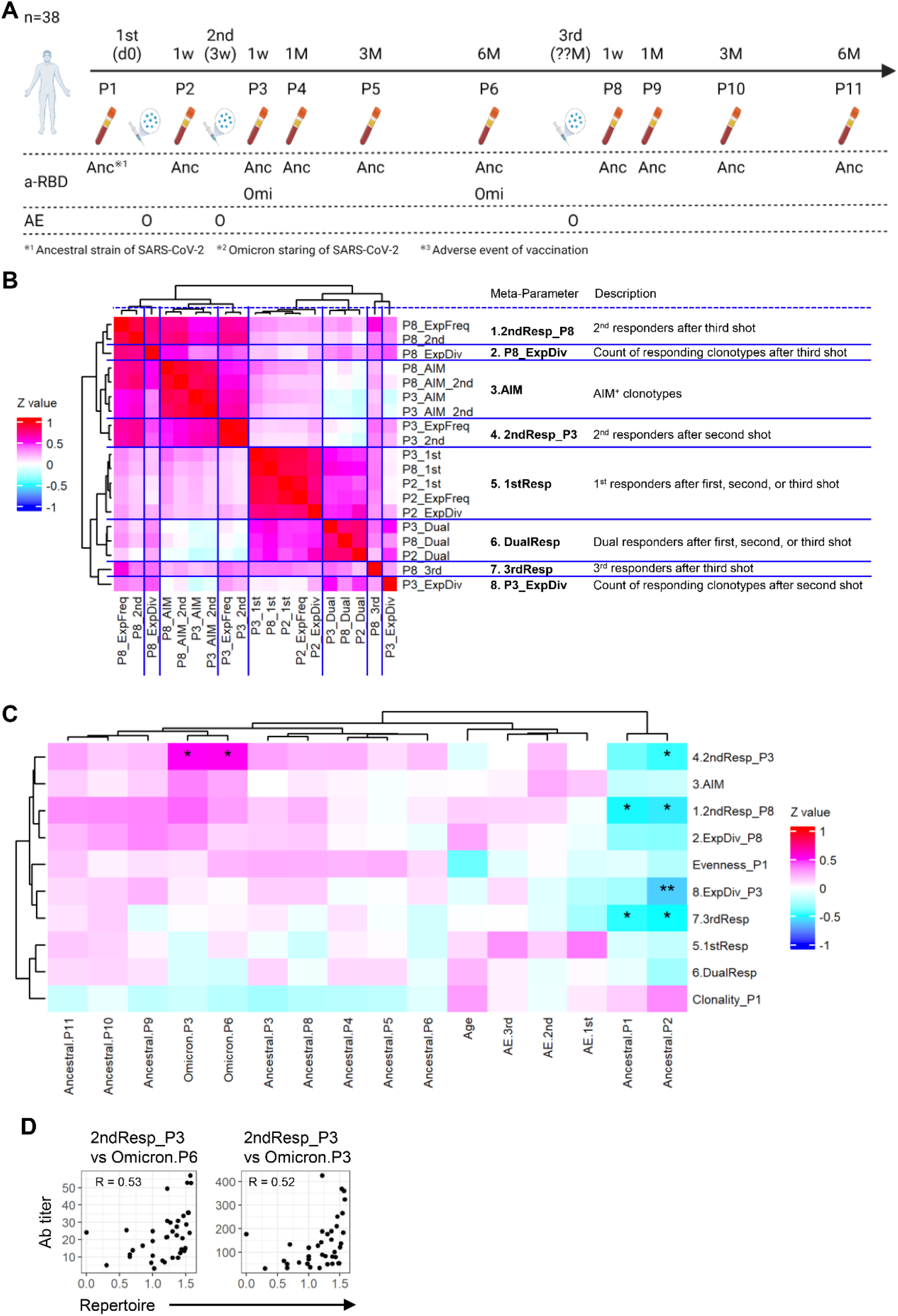
Correlation between CD8^+^ T cell repertoire meta-parameters and other variables regarding vaccination. (A) Schematic of time points of quantifying serum anti-RBD antibody titer and checking the presence of adverse vaccination events. Antibody titers against RBD of SARS-CoV-2 ancestral strain (“Anc”) and Omicron variant (“Omi”) were analyzed. (B) Heatmap of the correlation between CD8^+^ T cell repertoire parameters. The repertoire meta-parameter is calculated as the average of mutually correlated repertoire parameters (Materials and Methods). The names of repertoire meta-parameters and their descriptions are also shown. (C) Heatmap of the correlation between CD8^+^ T cell repertoire meta-parameters (row) and the age of participants, the titer of anti-SARS-CoV-2 RBD IgG antibodies (“Ancestral” or “Omicron”), or strength of the adverse events (“AE”) (column). (D) Correlation between the repertoire parameters (x-axis) and the IgG antibody titers to RBD of Omicron variant (“Omicron”) of SARS-CoV-2. The Spearman correlation coefficient between the two parameters is shown. (B and C). The Spearman correlation coefficient is colored in the heatmap. (C) q-values of the Spearman correlation coefficient between two parameters are plotted. **q<0.01; *q<0.05. (A) was created using BioRender.com.

### Temporal kinetics of T-cell clonotypes were analyzed within Spike epitopes

The results above demonstrate that the replacement of vaccine-responding clonotypes occurred between the first and the second shot. However, it remained unknown how this clonal replacement occurred (Figure 6A). One working hypothesis is “inter-epitope replacement,” in which different epitopes induce the 1^st^ and 2^nd^ responder clonotypes. The other hypothesis is “intra-epitope replacement,” in which both 1^st^ and 2^nd^ responder clonotypes exist in T cells specific to the same Spike epitope. To determine whether these two models explained the observed clonal replacement, we analyzed the temporal kinetics of Spike-specific T-cell clonotypes at an individual epitope level.

**Fig. 6.**
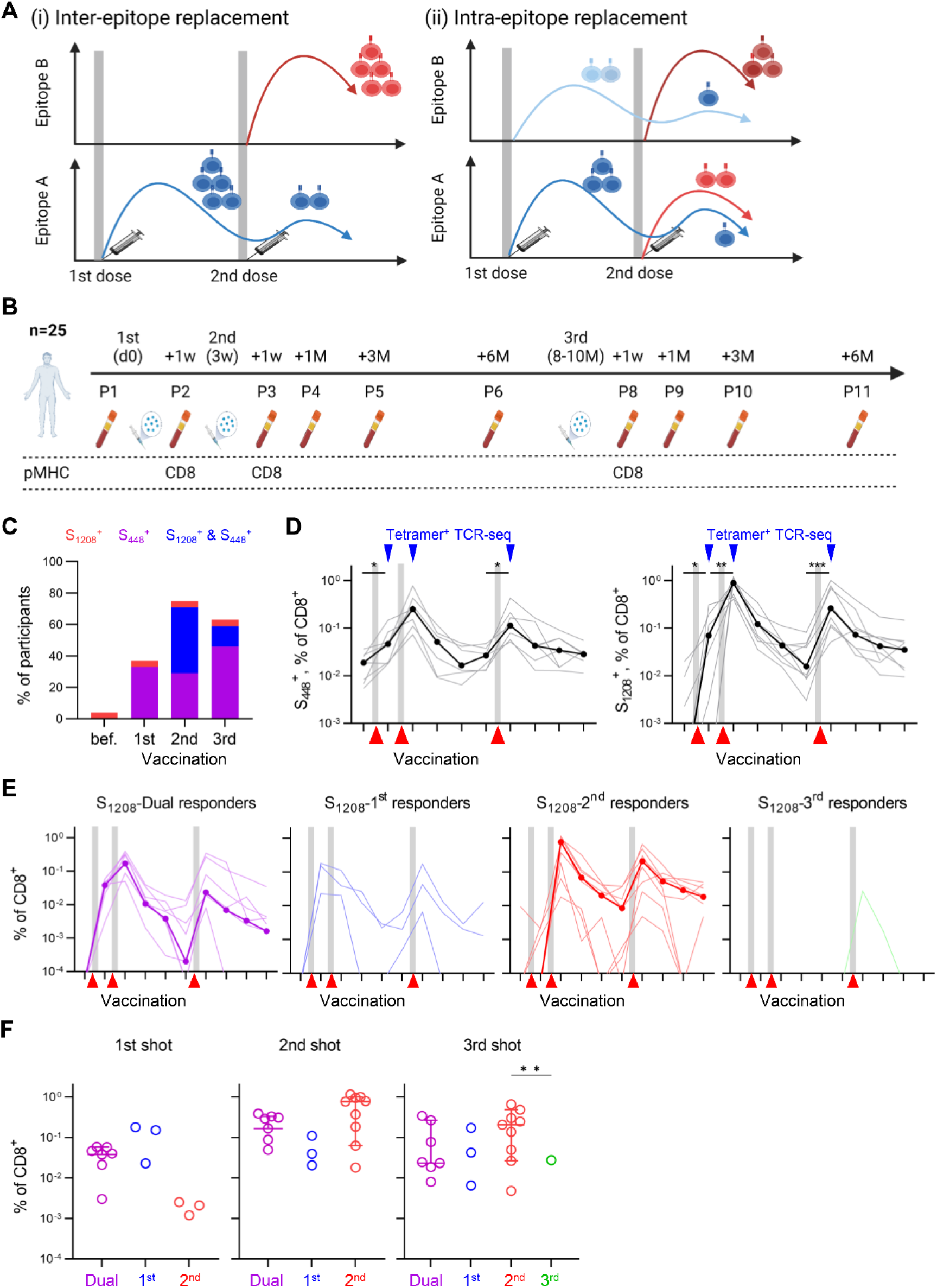
Analysis of Spike-specific clonotypes using pMHC-tetramer staining. (A) The working hypothesis for the clonal replacement during mRNA vaccination. (left) different epitopes induce the 1^st^ and 2^nd^ responder clonotypes (inter-epitope replacement). (right) Responding clonotypes are changed within each Spike epitope (intra-epitope replacement). (B) Schematic of time points of Spike peptide-MHC tetramer analysis. (C) Percentage of participants in whom S_448_ or S_1208_-specific T-cell responses were detected by flow cytometry. Spike-specific T-cell responses were defined as more than 0.1% tetramer^+^ cells among CD8^+^ T cells. (D) The total frequency of CD8^+^ S_448_^+^ (left) or S_1208_^+^ (right) clonotypes of participants in whom epitope-specific T-cell responses (“epitope-responsive participants”) were detected by repertoire analysis (thin line, grey). The median was also plotted (thick line, black). Epitope-responsive participants were defined in Supplementary Figure 6D. (E) Kinetics of S_1208_-specific 1^st^, 2^nd^, dual, and 3^rd^ responder clonotype frequencies. The thin line represents the total frequency for each participant. The thick line represents the median for S_1208_-responsive participants. (F) Comparison of the total frequency of S_1208_-specific 1^st^, 2^nd^, dual, and 3^rd^ responder clonotypes after each shot. The total frequency for 3^rd^ responders was plotted only after the third shot (right). n = 7 and 9 for S_448_- or S_1208_- responsive participants, respectively. (D and F) Friedman test with Dunn’s multiple comparisons was performed. ***P<0.001; **P<0.01; *P<0.05. (A and B) were created using BioRender.com.

For this purpose, two SARS-CoV-2 Spike epitopes were selected (S_448_ and S_1208_). These two epitopes are reported as immunodominant^32–34^ and presented by HLA-A24:02, the most frequent HLA class I allele in an Asian population. Then we prepared peptide-MHC (pMHC) tetramers and identified S_448_- or S_1208_-specific CD8^+^ T cells from A24:02^+^ participants using flow cytometry (Figure 6B and S6A). The positivity of S_448_-specific cells among CD8^+^ T cells increased after the first shot, while that of S_1208_-specific cells increased only after the second shot (Figure S6B). Moreover, the positivity of S_1208_-specific cells was significantly lower than that of S_448_-specific cells until after the first shot; however, they became comparable after the second shot (Figure S6C). Then, we performed TCR-seq on S_448_- or S_1208_-specific CD8+ T cells sorted from the samples after the vaccinations (P2, P3, and P8) and identified S_448_- or S_1208_-specific clonotypes (“Tetramer-TCRseq,” Figure S6D). Furthermore, for each participant, we determined whether the vaccine-induced S_448_- or S_1208_-specific T-cell responses were observed (S_448_- or S_1208_-responders, Figure S6D). Among nineteen participants in whom S_448_-specific clonotypes were detected, seven were classified as S_448_-responders, while twelve were not (Figure 6D and Figure S6E). All nine participants in whom S_1208_-specific clonotypes were detected were S_1208_-responders (Figure 6D). We focused on the S_448_- and S_1208_-responder participants (“respondent”) in the following section. In these respondents, the total frequency of S_448_- and S_1208_-specific clonotypes increased after the first shot and peaked after the second and third shots. At the peak of their response, the total frequency of S_1208_-specific clonotypes surpassed that of S_448_-specific ones (Figure S6F).

Finally, we examined whether the intra-epitope replacement of clonotypes was observed for S_448_ and S_1208_. S_448_- and S_1208_-specific clonotypes were classified into 1^st^, 2^nd^, dual, and 3^rd^ responders, and the total frequency of these responder clonotypes was calculated for each participant, as in Figure 1 (Figure 6E and Figure S6G). Among S_1208_-specific clonotypes, 2^nd^ and dual responders were observed in more than half of the respondents (Figure 6E, thick line). Moreover, the total frequency of S_1208_-specific dual responders tended to be higher than 2^nd^ responders after the first shot, while S_1208_-specific 2^nd^ responders tended to surpass the dual responders after the second shot (Figure 6F). These results are consistent with the replacement of clonotypes specific for the Spike epitope S_1208_. On the other hand, as for S_448_-specific clonotypes, only 2^nd^ responder clonotypes were observed in more than half of the respondents, while dual and 3^rd^ responders were rare, and 1^st^ responders were not observed (Figure S6G).

### Clonal replacement during mRNA vaccination occurs between and within Spike epitopes

TCRs recognizing the same pMHC often share conserved sequence motifs^35^. Based on this property, we identified clonotypes specific for Spike epitopes for which pMHC tetramer had been prepared (S_448_ and S_1208_), and also clonotypes specific for other Spike epitopes. From our original data of Tetramer TCR-seq and the public database of Spike-specific TCRs^36, 37^, shared TCR sequence motifs of Spike-specific clonotypes were constructed using tcrdist3 pipeline (Figure 7A)^38^. Then, we searched for T cell clonotypes possessing the Spike-specific TCR motifs, based on the participants’ HLA alleles (Figure 7A). Through this TCR motif analysis, we identified clonotypes specific for two Spike epitopes (S_269_-A02:01 and S_919_-B15:01), in addition to two epitopes used in Tetramer TCR-seq (S_448_-A24:02 and S_1208_-A24:02). Spike-specific clonotypes identified by TCR motif analysis were overlapped with those identified by Tetramer TCR-seq (Figure S7A), and there were no clonotypes to which different specificities were assigned between TCR motif analysis and Tetramer TCR-seq. On the other hand, the overlap between clonotypes with Spike-specific TCR motifs and AIM^+^ clonotypes was negligible (Figure S7B), suggesting that the Spike epitopes used in TCR motif analysis were presented to T cells less efficiently in our AIM assay with Spike peptide pool.

**Fig. 7.**
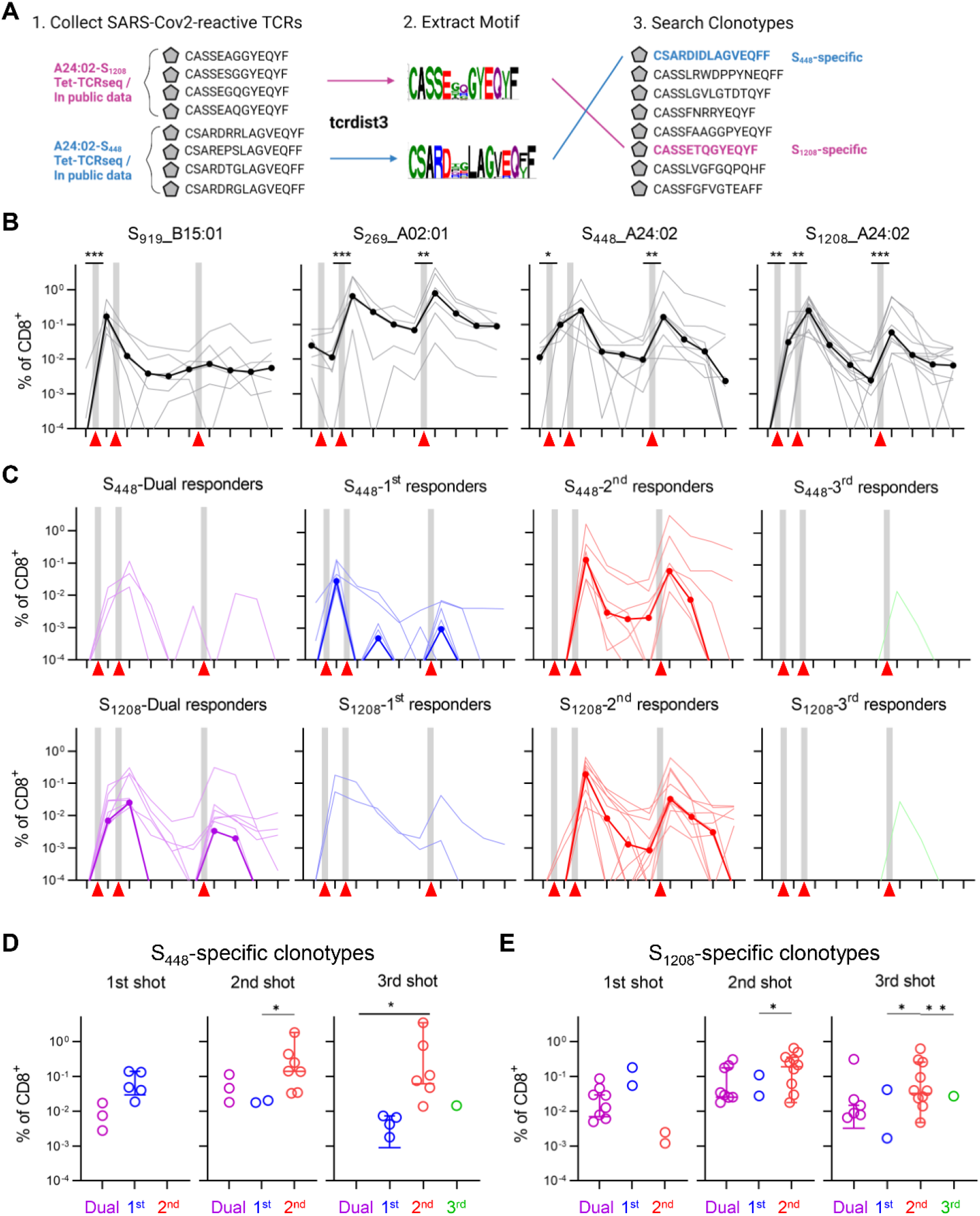
Analysis of Spike-specific clonotypes using TCR sequence motifs. (A) Schematic overview. (left) We used Spike-specific TCRs in three public databases and our original Tetramer-TCRseq datasets as input. (middle) Shared TCR sequence motifs of Spike-specific clonotypes were constructed using the tcrdist3 pipeline^38^. TCR sequence motifs are represented as logo plots, where the sizes and colors of the letters represent the frequency and characteristics of amino acids (red: acidic, blue: basic, green: polar, purple: neutral, black: hydrophobic). (right) T cell clonotypes possessing the Spike-specific TCR sequence motifs were searched. When searching clonotypes with sequence motif specific to each Spike-epitope, only participants with HLA that presents the epitope were analyzed. (B) The total frequency of CD8^+^ S_448_, S_1208_, S_269_, or S_919_-specific clonotypes of epitope-responsive participants (thin line, grey). The median was also plotted (thick line, black). Epitope-responsive participants were defined in Supplementary Figure 6D. (C) Kinetics of S_448_-(top) or S_1208_-(bottom) specific 1^st^, 2^nd^, dual, and 3^rd^ responder clonotype frequencies. The thin line represents the total frequency for each participant. The thick line represents the median for epitope-responsive participants. (D and E) Comparison of the total frequency of S_448_- (D) or S_1208_- (E) specific 1^st^, 2^nd^, dual, and 3^rd^ responder clonotypes after each vaccination. The total frequency for 3^rd^ responders was plotted only after the third shot. n = 8 (S_448_), 12 (S_1208_), or 6 (S_269_ and S_9_19). (B, D, and E) Friedman test with Dunn’s multiple comparisons. ***P<0.001; **P<0.01; *P<0.05. (A) was created using BioRender.com.

We then quantified the temporal kinetics of T-cell responses specific to the four Spike-epitopes by TCR motif analysis. Among these four Spike-epitopes, S_919_-B15:01 was most conserved between other human coronaviruses, while S_448_-A24:02 was least conserved and most unique to SARS-CoV-2 (Figure S7C). We observed that the temporal kinetics of the Spike-specific T-cell response differed between these epitopes (Figure 7B): S_448_- and S_1208_-specific responses were induced after the first shot and peaked after the second and third shots, as observed in Tetramer TCR-seq analysis. The S_269_-specific response also peaked after the second and third shots, while induction after the first shot was not observed (Figure 7B). The S_919_-specific response peaked after the first shot, but was not induced after the second and third shots (Figure 7B).

Furthermore, the total frequency of S_919_-specific clonotypes was significantly higher than S_1208_ and S_269_ after the first shot, while this was inverted after the second and third shots (Figure S7D). These results suggest that inter-epitope replacement of vaccine-responding clonotypes can occur between S_919_ and S_269_ in HLA-B15:01^+^ A02:01^+^ participants, and between S_919_ and S_448_ or S_1208_ in HLA-B15:01^+^ A24:02^+^ participants. However, we could not observe an example of the inter-epitope replacement within a single participant in our cohorts.

Furthermore, we classified Spike-specific clonotypes into 1^st^, 2^nd^, dual, and 3^rd^ responders to examine whether the intra-epitope replacement of clonotypes was observed. Among S_269_- and S_919_-specific clonotypes, only 2^nd^ and 1^st^ responders were observed in more than half of the respondents, respectively (Figure S6E and F). On the other hand, 1^st^ and 2^nd^ responders among S_448_-specific clonotypes, and 2^nd^ and dual responders among S_1208_-specific clonotypes were observed in more than half of the respondents (Figure 7C, thick line). As for S_448_-specific clonotypes, the total frequency of 1^st^ responders tended to be higher than 2^nd^ responders after the first shot, while 2^nd^ responders were significantly more abundant than 1^st^ responders after the second shot (Figure 7D). Similarly, as for S_1208_-specific clonotypes, the total frequency of dual responders tended to be higher than that of 2^nd^ responders after the first shot, while 2^nd^ responders tended to be more abundant than dual responders after the second shot (Figure 7E). Moreover, we observed replacement between 1^st^ and 2^nd^ responders in four of eight S_448_-respondents and between dual and 2^nd^ responders in three of twelve S_1208_-respondents (Figure S7H and G). These results provide examples of intra-epitope replacement of vaccine-responding clonotypes within Spike epitopes S_448_ and S_1208_. In summary, temporal tracking of Spike-specific clonotypes using TCR motif analysis demonstrated that the replacement of vaccine-responding clonotypes can occur within a Spike epitope, which does not require the differences in the target epitope between the 1^st^ and the 2^nd^ responder clonotypes.

## Discussion

Through a longitudinal TCR repertoire analysis combined with an AIM assay, single-cell TCR sequencing, and TCR motif analysis, we demonstrated that clonotypes newly expanded after the second vaccination largely replaced the clonotypes that had responded to the first shot. Moreover, although these 2^nd^ responder clonotypes expanded again with an effector memory phenotype, their diversity was further skewed after the third shot, and they were partially replaced by the mobilization of the 3^rd^ responders. Furthermore, this replacement of vaccine-responding clonotypes can occur not only between, but also within, clones specific for Spike epitopes. These data are consistent with a clonal replacement model of T-cell memory responses under repetitive antigen challenges in humans receiving SARS-CoV-2 mRNA vaccinations^18^.

To fully understand T cell clonal replacement following repeated mRNA vaccination, it is essential to clearly define the nature of 1^st^, 2^nd^, and dual responders. Secondary memory T cell responses are faster and more aggressive than the primary responses being medicated by naïve cells^39^. Based on this notion, we consider the 2^nd^ responder clonotypes as those that exerted the primary response after the first shot and recall response after the second shot. In our analysis, the primary response of most of the 2^nd^ responders was not detected after the first shot. We suppose this is because the cell count of 2^nd^ responders at the sampling point after the first shot is too small to detect through our bulk TCR-seq at P2, day 7 after the first shot.

In contrast, the dual and the 1^st^ responder clonotypes expanded quickly and vigorously after the first shot, which appeared closer to a recall response than a primary one.

Memory T cells cross-reactive to SARS-CoV-2 and ‘common cold’ coronaviruses (CCCoV) likely existed in many individuals before the COVID-19 pandemic^40–43^. Therefore, 1^st^ and dual responders may be “pre-existing memory” clonotypes that have been primed by CCCoV before the pandemic and exerted recall response by recognizing the SARS-CoV-2 Spike after the first shot. We further propose that their weak response after the second and third shot is that they differentiate into secondary memory with a weaker proliferative ability^44–46^. This is in line with the report that the contribution of CD4^+^ cross-reactive T cells to the Spike-reactive T-cell pool is minimal after vaccination against SARS-CoV-2^47^.

Consistent with this interpretation for 1^st^, 2^nd^, and dual responders, we propose that the T-cell response to S_919_, the most conserved Spike epitope analyzed in this study, was exerted by 1^st^ responder clonotypes. Moreover, the response to the less conserved S_269_ and S_1208_ epitopes was mainly exerted by 2^nd^ responders. On the other hand, the 1^st^ responder clonotypes were broadly observed in S_448_-specific clonotypes, contrary to the fact that S_448_ was least conserved among the four Spike epitopes, implying that pre-existing memory to SARS-CoV-2 S_448_ was less likely to exist. To explain this, we propose that there are pre-existing memory T cells that recognize epitopes similar to S_448_ derived from pathogens other than human coronavirus. Further studies are needed to prove the relationship between 1^st^ and dual-responder clonotypes and pre-existing memory T cells.

Determining whether clonal replacement through mRNA vaccination occurs between different Spike epitopes (inter-epitope replacement), or even within, a single Spike epitope (intra-epitope replacement) is an important step to understand the mechanism of clonal replacement. In the case of anti-Spike B cell responses following mRNA vaccination, differences in the response kinetics were reported between the target: the faster response to the S2 domain occurring after the first shot, which is cross-reactive to CCCoV, and the slower response to the S1 domain occurring after the second shot^48^. Like the B-cell response, our analysis identified a differential kinetics of Spike-specific T-cell responses between epitopes (i.e., response induced only after the first shot (S_919_), only after the second shot (S_269_), or both after the first and second shot (S_448_ and S_1208_)). This result supports the inter-epitope replacement model, possibly due to a shift of immunodominance in Spike epitopes through vaccination. Cases of inter-epitope replacement within a single participant were not observed, but might be found if we analyze larger cohorts with various combinations of HLA alleles, or if the Spike-specific TCR database was expanded to cover more diverse Spike epitopes. On the other hand, replacement between 2^nd^ responders and 1^st^ or dual responders was observed within the same epitopes (S_448_ and S_1208_) even in the same individual, which supports the inter-epitope replacement model. This observation implies the coexistence of two groups of clonotypes within a single Spike epitope: pre-existing memory clonotypes cross-reactive to CCCoV and newly expanded clonotypes specific to SARS-CoV-2. This complex nature of T-cell clonal responses within a single Spike epitope should be addressed through more intensive analysis of epitope-specific T cells and the identification of TCR motifs related to the responding patterns of clonotypes.

The third- and fourth-vaccination induces a more robust humoral and cellular immune response to SARS-CoV-2 than the second-vaccination^49, 50^, and increases the protection against SARS-CoV-2 infection, including the Omicron variant^51, 52^. As of April 2023, nearly 70 % and 50 % of the population in Japan have undergone more than three and four shots of vaccination, respectively. Our TCR repertoire analysis after the third vaccination demonstrated that the 2^nd^ responders were still dominant among vaccine-responding clonotypes and not significantly replaced by the 3^rd^ responders. However, after the third shot, 2^nd^ responders exhibited a more differentiated phenotype, and their diversity was significantly decreased compared to that after the second shot, resulting in the skewing of the 2^nd^ responders’ repertoire. The key question remains whether the diversity of 2^nd^ responders continued to decrease after four or more shots and whether 3^rd^ or 4^th^ responders can compensate for it. If this is the case, strategies may be needed to maintain the diversity of vaccine-responding clonotypes by replenishing the novel expanding ones under repetitive vaccination. Extending our TCR repertoire analysis after the fourth vaccination would provide critical information for optimizing the protocol for mRNA vaccination.

As far as we know, this study provided the first bulk TCR repertoire analysis dataset on more than 30 participants covering the entire time course of mRNA vaccinations (from the first shot to six months after the third shot). Moreover, the sampling time point is uniform across the participants, enabling the comprehensive analysis of the kinetics of T-cell clonotype expansion following vaccination. In addition to the analysis of the responding pattern of clonotypes, the comparison of the rate of contraction between vaccine-responding clonotypes would provide important information for understanding T-cell responses induced by mRNA vaccination. Moreover, the majority of the participants in our cohort reported neither clear infection history nor signs of past SARS-CoV-2 infection. Thus, our data reflect the effect of vaccination on T-cell repertoire more accurately. One defect in our dataset, however, is that the sampling point after the first vaccination (P2, day 7 after the first shot) was a few days earlier than the peak of T-cell responses to the first shot (around day 9-12)^53^. This earlier sampling can lead to an underestimation of the T cell response after the first shot: recall response of 1^st^ and dual responder clonotypes, and the primary response of 2^nd^ responders. However, an expansion of Spike-specific T-cell clonotypes (to S_919_, S_448_, and S_1208_) was observed at P2, confirming that this time point is not too early to detect vaccine-induced T-cell responses.

An important question is whether the clonal replacement we demonstrated in this study is specific to the mRNA vaccination or is common to the other types of vaccination or natural infections. We initially reported the clonal replacement in the antigen delivery systems using plasmid and viral vectors^18^. In addition, in the case of vaccination to yellow fever virus (YFV), the clonal composition of YFV-specific CD4^+^ T cells is reshaped by a single vaccination^54^, and the vaccine-responding CD8^+^ clones are also turned over between the prime and boost shots^55^. On the other hand, the trivalent influenza vaccination induces a response by T cells mainly recruited from the pre-existing memory clones^56^. This inconsistency in whether the clonal replacement occurs may be attributed to the type of vaccine (mRNA or live attenuated versus inactivated) or the novelty of the antigen (SARS-CoV-2 or YFV versus seasonal flu). Further study is needed to elucidate the requirements for clonal replacement after vaccination. In addition, considering that natural SARS-CoV-2 infection generates tissue-resident memory T cells (T_RM_) in the lung^57^, the replacement of antigen-specific clonotypes in blood after re-infection, if it occurs, may be associated with the re-distribution of older clonotypes from the blood to peripheral tissues and their differentiation to T_RM_. Collectively, along with the temporal analysis of blood T cell repertoire as done in the current study, a systemic analysis of the T cells in peripheral tissues is essential to understand the kinetics of T-cell clonal response by natural infection.

In summary, we demonstrated the replacement of memory T-cell clones during SARS-CoV-2 mRNA vaccination. In contrast to humoral immunity mediated by memory B cells and long-lived plasma cells, where the memory response to the first viral variant encountered in childhood is preferentially employed even after the challenge of other variants in adults (original antigenic sin)^58, 59^, T cells appear to be able to reshape the memory population after repetitive challenges. This “re-writability“ of T cell immunological memory may enable them to effectively generate responses to variant strains of the virus.

## Methods

### Study design

38 healthcare workers (≥20 years old) at Nara Medical University in Japan who had received the first shot of Pfizer-BNT162b2 vaccine between 10 March and 30 March 2021 were invited to participate as volunteers in the study. The Nara Medical University Ethics Committee approved the study (No. 2954 and 3168). In this cohort, no one reported a SARS-CoV-2 infection history before or during the initial sampling period. Four and two participants showed signs of past SARS-CoV-2 infection before the first and third vaccination, respectively (inferred by the anti-Nucleoprotein (N) antibody titer). Peripheral blood samples were collected before receiving the first dose (P1), 1 week after the first shot (P2), 1 week (P3), 1 month (P4), 3 months (P5), and 6 months (P6) after the second shot, and 1 week (P8), 1 month (P9), 3 months (P10), and 6 months (P11) after the third shot. One participant (ID18) was not sampled at P11 because they had received the fourth vaccination before this time point. Participants were asked to report the following adverse events after each vaccination: pain, swelling, heat, fever, fatigue, headache, chills, joint pain, and muscle pain. PBMCs were isolated from whole blood with BD vacutainer CPT tubes (BD reagents used in this study are summarized in Supplementary Table S3), resuspended in CELLBANKER 1 (Takara), and stored at -80°C. The detailed study design was described in Kitabatake et al.^19^.

### Study approval

The study was conducted in accordance with the Declaration of Helsinki and Good Clinical Practice Guidelines, following approval by the ethics board in each institution. All study participants provided written informed consent.

### Evaluation of anti-SARS-CoV-2 antibodies

Serum samples were tested for antibody titer against RBD of SARS-CoV-2 ancestral strain, SARS-CoV-2 Omicron spike variant protein, and nucleocapsid protein of SARS-CoV-2 ancestral strain, were tested using RayBio COVID-19 S1 RBD protein Human IgG ELISA Kit (RayBiotech), RayBio COVID-19 Omicron Spike Variant Human IgG ELISA Kit (RayBiotech), and RayBio COVID-19 N protein Human IgG ELISA Kit (RayBiotech) per the manufacturer’s instructions, respectively.

### T-cell isolation from PBMCs

CD4^+^ and CD8^+^ T cells were magnetically isolated from PBMC. PBMC were thawed by warming frozen cryovials in a 37°C water bath. Cells were stained for 20 minutes at 4°C with 1^st^ antibody mix (Antibody Mix 1 in Supplementary Table S4). After washing with MACS buffer [0.5% BSA (Nacalai), 2 mmol/L EDTA (NIPPON GENE) in PBS (Nacalai)], Cells were stained for 20 minutes at 4°C with 2^nd^ antibody mix; BD IMag anti-PE Magnetic Particles-DM (Clone E31-1459, BD) and anti-APC-Biotin (clone APC003, BioLegend). Then, the cell suspension was diluted with MACS buffer and set on a magnet plate (DynaMag96SideSkirted, Veritas). The positive fraction was collected as CD4^+^ T cells. The negative fraction was stained with BD IMag Streptavidin Particles Plus-DM (BD) for 20 minutes at 4°C. Then, the cell suspension was diluted with MACS buffer and set on a magnet plate. The positive fraction was collected as CD8^+^ T cells and B cells. The counts of CD4^+^ T cells, CD8^+^ T cells, and B cells were determined using Flow-Count fluorospheres (Beckman) and a Cytoflex flow cytometer (Beckman) and FlowJo software (version 10.8.1; BD Biosciences, software and algorithms used in this study are summarized in Supplementary Table S5).

### HLA genotyping

Genome DNA was extracted from PBMC using Geno Plus Genomic DNA Extraction Miniprep System (VIOGENE) using serum and animal cell protocol with RNase A treatment. DNA was quantified on the NanoDrop (Thermo Fisher Scientific). HLA genotyping was performed on genome DNA using the reverse sequence-specific oligonucleotide probe Luminex method at HLA Laboratory (Kyoto, Japan).

### AIM assay and sorting Spike-reactive T cells

AIM assay was performed on PBMC samples collected at P2, P3, and P8. PBMC were thawed by warming frozen cryovials in a 37°C water bath and resuspended in 10mL of in Culture Medium (CM) [10% fetal bovine serum (MP Bio Japan), 1% HEPES (Nacarai) and 1% penicillin/streptomycin (Nacarai) in RPMI-1640 medium (Nacarai)]. Cells were resuspended in 1mL of CM, then plated in 24-well plates and incubated overnight (about 16 hours) in an incubator at 37°C, with 5% CO_2_. After incubation, cells were then stimulated for 24 hours in stimulation medium [CM with BD FastImmune CD28/CD49d Costimulatory Reagent (final x100, BD Biosciences) and anti-human CD40 (final concentration of 0.5μg/mL, clone HB14, BioLegend)] with or without PepTivator SARS-CoV-2 Prot_S Complete (final concentration of 1 mg/mL, Miltenyi). 20 hours post-stimulation, an anti-CD107a-FITC antibody (final concentration of 5μg/mL, clone H4A3, BioLegend) was added to the culture. After four hours, cells were washed in PBS supplemented with 10% FBS (10% FBS/PBS). Cells were stained for 20 minutes at 4°C with antibody mix 2 (Supplementary Table S4). Cells were washed once in 10% FBS/PBS. AIM^+^ non-naïve CD4^+^, AIM^-^ non-naïve CD4^+^, AIM^+^ non-naïve CD8^+^, and AIM^-^ non-naive CD8^+^ cells were sorted using FACS Aria II or Aria III (BD Biosciences) and FACSDiva software (BD Biosciences, v8.0.1). AIM^+^ CD4^+^ T cells were defined by dual expression of CD69 and CD40L. AIM^+^ CD8^+^ T cells were defined by expression of either CD107a or CD137. Nonviable cells were excluded from the analysis based on forward and side scatter profiles and propidium iodide staining (PI, Sigma-Aldrich). Data were analyzed using FlowJo software.

### Sample preparation for single-cell analysis

PBMC thawed as described above were resuspended with Cell Staining Buffer (BioLegend) supplemented with Human TruStain FcX. After 2-3 minutes, 2x AbSeq master mix of Immune Discovery Panel (BD Biosciences) prepared as the provider’s protocol was added to the cell suspension and stained for 15 minutes on ice. Then, antibody mix 3 (Supplementary Table S4) with Sample Tag oligonucleotide-conjugated antibodies (Single-Cell Multiplexing Kit, BD Biosciences) was added to the cell suspension and left on ice for 15 minutes for staining. The correspondence between samples and Sample Tags used was described in Supplementary Table S2. Stained cells were washed with Cell Staining Buffer. Then non-naive CD4^+^ and CD8^+^ cells (ID2 and ID27) or only non-naive CD8^+^ cells (ID10, ID19, ID22, ID31, and ID35) were sorted using FACS Aria II or Aria III and FACSDiva software (v8.0.1). Nonviable cells were excluded from the analysis based on forward and side scatter profiles and PI staining. Data were analyzed using FlowJo software (version 10.8.1).

### Tetramer staining and sorting Spike-specific T cells

S_448_ and S_1208_ peptides were purchased from GenScript. Peptides-MHC class I tetramers for SARS-CoV-2 Spike epitopes (S_448_ and S_1208_) were generated by QuickSwitchTM Quant HLA-A*24:02 Tetramer Kit-BV421, and -PE (MBL), respectively, according to the manufacturer’s protocol. The rate of peptide exchange was reported to be more than 95% in our previous report^32^. Thawed PBMCs were treated with a protein kinase inhibitor, Dasatinib (50 nM, AdooQ), for 30 min at 37 °C, and Human TruStain FcX for 5 minutes at room temperature. Then, PBMCs were stained with tetramers for 30 minutes at room temperature. After tetramer staining, cells were stained for antibody mix 4 (Supplementary Table S4) for 20 minutes at room temperature. After washing twice with FACS buffer, CD8^+^ tetramer^+^ cells were sorted using FACS Aria II or Aria III and FACSDiva software (v8.0.1). Nonviable cells were excluded from the analysis based on forward and side scatter profiles and PI staining. Data were analyzed using FlowJo software (v10.8.1).

### Library preparation and sequencing for TCR repertoire analysis

TCR sequencing libraries for next-generation sequencing were prepared according to the previous report^60^. In brief, mRNA in T-cell lysates were captured by incubating with 10 μL of Dynabeads M270 streptavidin (Thermo Fisher Scientific) bound to 2.5 pmol of BioEcoP-dT25-adapter primers for 30 minutes at room temperature under gentle rotation. mRNA-trapped beads were suspended in 10 µL of RT mix and incubated for 60 min at 42°C to perform reverse transcription (RT) and template-switching. The composition of the RT mix was as follows; 1× First Strand buffer (Thermo Fisher Scientific), 1 mM dNTP (Roche), 2.5 mM DTT (Thermo Fisher Scientific), 1 M betaine (Sigma-Aldrich), 9 mM MgCl2 (NIPPON GENE), 1 U/µL RNaseIn Plus RNase Inhibitor (Promega), 10 U/µL Superscript II (Thermo Fisher Scientific), and 1 µM of i5-TSO. To amplify the TCR cDNA containing complementarity determining region 3 (CDR3), nested PCR of the TCR locus was performed using KAPA Hifi Hotstart ReadyMix (KAPA Biosystems), followed by the purification of PCR product using an AM Pure XP kit (Beckman Coulter) at a 0.7:1 beads-to-sample ratio. The third PCR was performed to amplify TCR libraries and add adaptor sequences for the next-generation sequencer. The third-PCR products were purified as described for the second PCR. Final TCR libraries, whose lengths were about 600 base pairs, were quantified using a KAPA SYBR Fast qPCR Kit (KAPA Biosystems), and size distribution was analyzed by Microchip Electrophoresis System MultiNA (Shimadzu). TCR libraries were sequenced using an Illumina Novaseq 6000 S4 flowcell (67 bp read 1 and 140 bp read 2) (Illumina) using NovaSeq 6000 S4 Reagent Kit v1.5 (200 cycles, Illumina). Only read2 contained the sequence regarding the definition of T-cell clones. Oligonucleotides and reaction conditions for library preparation were summarized in Table S6 and S7, respectively.

For tetramer TCR-seq, the libraries were prepared on samples with (i) more than 100 tetramer^+^ cells and (ii) the proportion of tetramer^+^ cells was more than 0.1% in CD8^+^ T cells. S_1208_-tetramer^+^ samples with a high background (ID22) were excluded from the analysis.

### Library preparation and sequencing for single-cell analysis

Single-cell sequencing libraries were prepared according to the previous report^60^. Live cells were stained with Calcein AM (Nacalai) and counted using Flow-Count fluorospheres and a Cytoflex flow cytometer. About 25,000 - 30,000 labeled cells were loaded on a BD Rhapsody cartridge (BD) and processed using BD Rhapsody™ Cartridge Reagent Kit (BD) following the manufacturer’s instructions. Single-cell mRNA, AbSeq barcodes, and Sample Tag barcodes were captured by BD Rhapsody beads coated with poly(T) oligonucleotide with a unique cell barcode and molecular barcode on each bead. Single-cell cDNA synthesis was performed using BD Rhapsody cDNA Kit (BD) following the manufacturer’s protocol. Targeted mRNA, TCRseq, and Abseq Index libraries were prepared using Immune Response Panel HS (BD) according to the manufacturer’s instructions with the following modifications. (1) We used the original set of primers for TCR amplification, as summarized in Table S6. (2) We performed additional PCRs to introduce a unique dual index for sequencing (3rd PCR for mRNA and TCR), as described in Table S8. (3) We used the ProNex Size-Selection DNA Purification System (Promega) to purify PCR products, as summarized in Table S8. Amplified libraries were quantified using a KAPA SYBR Fast qPCR Kit (KAPA Biosystems), and size distribution was analyzed by Microchip Electrophoresis System MultiNA (Shimadzu). Targeted RNAseq, TCRseq, and AbSeq libraries were sequenced on an Illumina Novaseq 6000 S4 flowcell (67 bp read 1 and 140 bp read 2) using NovaSeq 6000 S4 Reagent Kit v1.5 to a depth of approximately 20,000 reads, 7,000 reads, and 10,000 reads per cell, respectively.

### Data processing of TCR sequencing reads

Data processing of TCRseq was performed as previously reported^60^. Briefly, Adapter trimming and quality filtering of sequencing data were performed using Cutadapt-3.2^61^ and PRINSEQ-0.20.4^62^. Sequencing data were processed by MiXCR-3.0.5^63^. In MiXCR, filtered reads were aligned to reference mouse TCR V/D/J sequences registered in the international ImMunoGeneTics (IMGT) information system. The parameters used in MiXCR are summarized in Supplementary Table S9. The Variable (V) and Joining (J) segments of TCRs were represented in IMGT gene nomenclature. T-cell clones were determined as TCR reads with the same TCR V segment, J segment, and CDR3 nucleotide sequence.

For bulk TCRseq data, we regarded the CD4^+^-CD8^+^ overlapping clones in the same participant as contamination during T-cell isolation from PBMC because CD4^+^ and CD8^+^ T-cell repertoire are mutually exclusive^64^. Thus, these overlapping clones were categorized into either CD4^+^ or CD8^+^ according to their dominance in CD4^+^ and CD8^+^ repertoire using VDJtools-1.2.1^65^. Inter-sample contamination within samples using the same UDIs was excluded by the “Decontaminate“ function of vdjtools with the parameter “-r 10“. The sequencing reads of bulk TCR-seq samples were normalized to 100,000 reads by the “DownSample” command of VDJtools-1.2.1^65^. To preserve the minor clonotypes, we performed the normalization ten times and pooled these down-sampled repertoire data for each sample by the “JoinSamples” command of VDJtools-1.2.1. The sequencing reads of AIM TCR-seq or tetramer TCR-seq samples were normalized to 16 times the cell count (AIM assay-TCRseq data) in each sample by the “DownSample” command of VDJtools-1.2.1^65^, and TCR clones with 15 or fewer reads were regarded as sampling noise and removed from the dataset^64^.

### Identifying vaccine-responding clonotype using bulk TCR-seq dataset

Clonotypes that significantly expanded after vaccination (vaccine-responding clonotype) were identified by differential abundance analysis using the beta-binomial model^20^. Before performing differential abundance analysis on clonotype frequency, the total count of bulk TCRseq data was adjusted to 100,000. Then, vaccine-responding and vaccine-increased clonotypes are defined for each vaccination (P2, P3, or P8).

Clonotypes are regarded as “responded to vaccination” when (i) they had a significant increase in their frequency and (ii) their frequency was increased more than two-fold between before and after the vaccination. Vaccine-responding clonotypes were also identified between the baseline (P0) and after the second shot (P3). In addition, clonotypes are regarded as “increased after vaccination” when (i) their frequency was increased between before and after the vaccination, (ii) their frequency was increased more than two-fold between the baseline (P1) and after the vaccination, and (iii) their frequency was more than 0.0025% after the vaccination. Finally, Clonotypes that did not meet the requirements for vaccine-responding or vaccine-increased clonotypes were regarded as “did not respond to vaccination.” The beta-binomial model incorporates time-dependent variance of clonotype frequency using normal peripheral blood T cell repertoire variance over two weeks^20^.

### Identifying 1^st^, 2^nd^, dual, and 3^rd^ responder clonotypes in bulk TCR-seq dataset

1^st^ responder clonotypes were defined as clonotypes that responded to the first shot, but did not respond to the second one. 2^nd^ responder clonotypes were defined as clonotypes that responded to the second shot while did not respond to the first shot. Dual responder clonotypes were defined as clonotypes that responded or increased after the first and second shots, and that responded between the baseline (P0) and after the second shot (P3). 3^rd^ responder clonotypes were defined as clonotypes that were not detected before the third shot and responded to the third shot. We excluded clonotypes whose frequency was over 0.01% at baseline from 1^st^, 2^nd^, and dual responders.

### Identifying Spike-reactive clonotypes through AIM TCR-seq dataset

Spike-reactive clonotypes were identified using the dataset of AIM TCR-seq as in Figure 2B. “AIM^+^ enrichment ratio” was calculated for each clonotype as (frequency in AIM^+^ repertoire + 5×10^-6^) / (frequency in AIM^-^ repertoire + 5×10^-6^). Clonotypes with an AIM^+^ enrichment ratio over 16 were defined as Spike-reactive clonotypes, and their frequency was tracked in the bulk TCR-seq dataset.

### Calculation of clonality or evenness of TCR repertoire

The 1 - Pielou index was used to evaluate the clonality of TCR repertoire, which was calculated using the formula 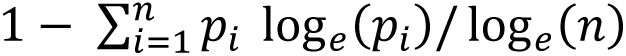, where *p_i_* is the frequency of clone *i* for a sample with *n* unique clones. The DE90 index was used to evaluate the evenness of the TCR repertoire, which was calculated as the ratio between the number of clonotypes whose cumulative frequency is 90% and the total number of clonotypes in the repertoire^66^.

### Identifying Spike-specific clonotypes through tetramer TCR-seq dataset

S_448_- and S_1208_-specific clonotypes were identified using the dataset of tetramer TCR-seq and bulk TCR-seq as in Figure S6D. “Tetramer^+^ enrichment ratio” was calculated for each clonotype as (frequency in tetramer^+^ repertoire + 5×10^-6^) / (maximum frequency in bulk TCR-seq datasets from P1 to P11 + 5×10^-6^). Clonotypes with a tetramer^+^ enrichment ratio over 16 were defined as S_448_- or S_1208_-specific clonotypes, and their frequency was tracked in the bulk TCR-seq dataset. Then, epitope-responsive participants were defined as those whose total frequency of epitope-specific clonotypes was (i) more than 0.05% and (ii) 5-fold increased after the first or the second shot (red).

### Identifying Spike-specific clonotypes through TCR motif analysis using tcrdist3

To identify Spike-specific clonotypes through TCR motif analysis in our dataset, we utilized the framework introduced in Mayer-Blackwell et al.^38^. In this framework, we first formed meta-clonotypes (groups of TCRs that were biochemically similar) of Spike-specific TCRs. Then we searched TCRs conformant to the definition of the meta-clonotypes (meta-clonotype conformant TCRs).

Spike-specific meta-clonotypes are formed using Spike-specific TCR sequences from two public databases (vdjdb and immune epitope database: IEDB) and our tetramer TCR-seq datasets as an input. In this step, the TCR sequence motif and optimal radii were estimated for each meta-clonotype at which background TCRs were detected at a frequency of < 10^-5^. Then, Spike-specific TCRs were searched in our dataset using these Spike-specific meta-clonotypes and meta-clonotypes reported in Mayer-Blackwell et al.^38^, which were identified from MIRA Spike antigen stimulation assay dataset^37^ with significant association for HLA-restricted epitopes. In this step, the query list of Spike-specific meta-clonotypes was generated for each participant by selecting meta-clonotypes whose epitope-presenting HLA allele matched the participant’s HLA genotypes. The frequency of Spike-specific clonotype was tracked in bulk TCR-seq dataset, and epitope-responsive participants were defined as for the tetramer TCR-seq data analysis.

### Data processing of scRNAseq reads

Data processing of scRNAseq reads was performed as described in Shichino et al ^67^. Adapter trimming of sequencing data was performed by using cutadapt 3.4^61^. Filtered reads were chunked into 16 parts for parallel processing using Seqkit 0.15.0^68^. Filtered cell barcode reads were annotated by Python script provided by BD with minor modifications to be compatible with Python3.8. Associated cDNA reads were mapped to reference RNA (build GRCh38 release-104) using bowtie2-2.4.2^69^ by the following parameters: -p 2 -D 20 -R 3 -N 0 -L 8 -I S,1,0.75 –norc –seed 656565 –reorder. Then, cell barcode information of each read was added to the bowtie2-mapped BAM files, and read counts of each gene in each cell barcode were counted by using mawk. The resulting count data was converted to a genes x cells matrix file, and the knee-plot inflection threshold was detected using the DropletUtils package in R version 4.1.2^70^. In addition, we further estimate background beads using the emptyDrops formula in the DropletUtils package. Cells of total read count over inflection point were considered valid cells.

### Data processing of sample tag reads

For sample tag data, adapter trimming of sequencing data was performed using cutadapt 3.4^61^. Filtered reads were chunked to 64 parts for parallel processing using Seqkit 0.15.0^68^. Filtered cell barcode reads were annotated using the Python script provided by BD with a minor modification to allow compatibility with Python3.8. Associated sample tag reads were mapped to known barcode fasta using bowtie2-2.4.2^69^ with the following parameters: -p 2 -D 50 -R 20 -N 0 -L 14 -i S,1,0.75 –norc –seed 656565 –reorder –trim-to 3:40 –score-min L,-9,0 –mp 3,3 –np 3 –rdg 3,3.

The cell barcode information for each read was added to the bowtie2-mapped BAM files, and read counts of each Tag in each cell barcode were counted using mawk. The resulting count data were converted to a Tags x cells matrix file using the data.table package in R 3.6.3, and the top 1M cell barcodes were extracted. For the assignment of individual tags to cell barcodes, read counts of each tag in each valid cell barcode, defined by the cDNA matrix, were extracted from the tag/cell barcode expression matrix. Unassigned cell barcodes were labeled as “not-detected” cells. The sum of the total read counts of each tag was normalized to the minimum sum count of each tags, and log_2_ fold-change between first most tag counts and second most tag counts within each cell barcode. Each cell barcode was ranked by the fold-change ascending order, and top N cells were identified as doublets (N was calculated as theoretically detectable doublets calculated using the Poisson’s distribution based on the number of loaded cells, total Rhapsody well number, and the number of tags used).

### Data processing of AbSeq reads

For AbSeq data, adapter trimming of sequencing data was performed using cutadapt 3.4^61^. Filtered reads were chunked into 64 parts for parallel processing using Seqkit 0.15.0^68^. Filtered cell barcode reads were annotated by Python script provided by BD with minor modifications to be compatible with Python3.8. Associated AbSeq reads were mapped to known barcode fasta by using bowtie2-2.4.2^69^ by the following parameters: - p 2 -D 20 -R 3 -N 0 -L 8 -I S,1,0.75 –norc –seed 656565 –reorder–trim-to 3:39 –score-min L,-9,0 –mp 3,3 –np 3 –rdg 3,3. Then, the cell barcode information of each read was added to the bowtie2-mapped BAM files, and read counts of each AbSeq tag in each cell barcode were counted using mawk. The resulting count data were converted to tags x cells matrix file by using the data.table package in R version 4.1.2, and top 1M cell barcodes were extracted.

### Correct batch effect in scRNAseq and AbSeq data

scRNAseq/AbSeq data analysis was performed using the R software package Seurat v4.0.1^71^. Expression data of targeted panel genes and surface proteins were converted to the Seurat object. CD4^-^ CD8^+^ cells were extracted based on the expression level of CD4 and CD8 proteins determined by AbSeq. Then, expression data was log-normalized and scaled using NormalizeData and ScaleData functions, respectively. For scRNAseq data, the batch effect was corrected as in Stuart et. al.^72^, using reciprocal principal component analysis (PCA) in Seurat. Then, PCA was performed using RunPCA (the number of calculated principal components (PCs) was 50). The enrichment of each PC was calculated using the JackStraw and ScoreJackStraw function (num.replicate = 100), and PCs that were significantly enriched (p ≤ 0.05) were selected for further analysis. For AbSeq data, genes unrelated to the phenotype of CD8^+^ T cells (CD14, CD16, CD19, CD3, CD4, CD8, CXCR5, IgD, and IgM) were excluded, then, the batch effect of selected genes (CD27, CD28, CD56, CD62L, and CD161) were corrected using the ADTnorm function in the package ADTnorm^73^. The expression data of the other genes (CCR7, CD11c, CD25, CD45RA, CD127, CD134, CD137, CD183, CD196, CD272, CD278, CD279, CXCR6, GITR, HLA-DR, and Tim3) was log-normalized using NormalizeData function. After that, data scaling and PCR were performed using ScaleData and RunPCA function.

### Cell clustering through integrative analysis of scRNAseq and AbSeq data

Cell clustering was performed based on the weighted-nearest neighbor graph using FindMultiModalNeighbors and FindClusters against the significant PCs of scRNAseq and AbSeq^74^. The dimensional reduction was performed using RunUMAP based on the nearest neighbor graph determined by FindMultiModalNeighbors. Marker genes of each identified cluster were identified using the FindAllMarkers function (test.use=“Wilcox“, min.pct=0.1, return.thresh=0.05). Clusters constituted with contaminant cells determined by marker genes were excluded, and the above analysis procedure was repeated.

### Transcriptional signature analysis in scRNAseq

We computed the extent to which gene signatures were expressed in cells using Seurat’s “AddModuleScore” function. Signature genes related to the T-cell phenotypes were obtained from the dataset published by Galleti et al.^75^. Effector and Cell cycle signature genes were obtained based on the descriptions of Immune Response Panel HS. Genes used for calculating signature scores are summarized in Supplementary Table S10.

### Data processing and analysis of scTCRseq

Raw data from single-cell TCR sequencings were processed by the pipeline produced by ImmunoGeneTeqs, Inc., as described previously^60^. Briefly, sequencing reads were separated by cell barcode, then adapter trimming and quality filtering of sequencing data were performed using cutadapt 3.4^61^. PCR and sequencing errors were corrected by lighter v1.1.2^76^ with the following parameters: -newQual 25 -maxcor 4 -K 20. Error-corrected sequence reads were processed by MiXCR^63^ to generate the list of TCRα or TCRβ sequences for each cell barcode, using reference human TCR V/D/J sequences registered in the international IMGT information system. Quality filtering of cell barcodes was performed with the following criteria: (1) more than 32 TCR reads were detected. (2) the proportion of the most frequent TCR sequence in cell barcodes was over 0.6. (3) the cell barcode was annotated to the T cell cluster in scRNAseq/AbSeq analysis. Parameters used in MiXCR are summarized in Supplementary Table S11. Finally, the most frequent sequence of TCRα and TCRβ was adopted and paired in each cell barcode. Identified TCRα or TCRβ sequence was imported into the meta.data matrix in the Seurat object of scRNAseq/AbSeq data based on the cellular barcode identities. T-cell clones were determined as cells with the same TCRβ sequence defined by the V segment, J segment, and CDR3 nucleotide sequence. The frequency of each clone was calculated as the proportion of cells belonging to the clone in all T cells whose TCRβ sequence was assigned.

### Calculating the TCR repertoire meta-parameter for correlation analysis

Co-regulated TCR repertoire parameters (“repertoire meta-parameters“) were independently determined for CD4^+^ and CD8^+^ T cells. First, a Spearman correlation coefficient was calculated for all pairs of repertoire parameters; then, repertoire parameters were grouped into 8 clusters based on the matrix of correlation coefficient values. TCR repertoire meta-parameter were calculated by averaging the rank of parameters in each cluster. Meta-parameters were used for correlation analysis.

### Quantification and statistical analysis

Statistical analyses were performed using GraphPad Prism (ver9.5.1) software (GraphPad Software). Friedman test with Dunn’s multiple comparisons test was performed on the comparison between the responding patterns of clonotypes or temporal comparison of clonotype frequency. Wilcoxon matched-pairs signed rank test was performed on the comparison of 2^nd^ responder clonotypes after the second and the third shot or comparison of the frequency of S_448_ and S_1208_ tetramer^+^ cells determined by flow cytometry. The Spearman test was performed on the correlation analysis between repertoire meta-parameters and clinical parameters. Mann-Whitney test was performed on the comparison of the total frequency of S_448_- and S_1208_-specific clonotypes determined by tetramer TCR-seq in responding participants. Kruskal-Wallis test with Dunn’s multiple comparisons was performed on the comparison of the total frequency of S_448_-, S_1208_-, S_269_-, and S_919_-specific clonotypes determined by TCR motif analysis in responding participants. q-values are estimated from simultaneous testing (Figure 3H and Figure 5, Supplementary Figure S5) using “qvalue” packaged in R^77^. Asterisks to indicate significance corresponding to the following: ***P<0.001; **P<0.005; *P<0.05; n.s. non-significant (P≥0.05).

### Data and materials availability

All sequencing data (Bulk TCRseq, and Single-cell RNA/TCR sequencing) have been deposited at GEO and are publicly available as of the date of publication (GEO: GSE210231). All original code has been deposited at Github (https://github.com/hiro-aoki-mriid/Covid_TCRrepertoire). The list of reagents, antibodies, and algorithms used in this study was summarized in Supplementary Tables S3, 4, and 5, respectively.

## Acknowledgments

This work was supported by the Japan Society for the Promotion of Science (JSPS) under Grant Numbers 17929397, 20281832, and 22H05064, and Japan Agency for Medical Research and Development (AMED) under Grant Number JP22fk0310509, JP22ama221306, and P21gm6210025. We want to thank Y. Hara for advice in cell sorting, members of IGT. Inc. for expert technical assistance in TCR sequencing, M. Takahashi, D. Komura, and S. Ishikawa for advice in data analysis. We thank Dr. David L. Woodland for excellent English language editing. Fig. 1A, 1E, 2B, 4E, 6A, and 7A were created with BioRender.com.

## Author Contributions

HA, MK, TI, and SU performed conceptualization. HA performed data curation, formal analysis, and visualization. HA and SS developed software. HA, SS, AH, CM, and SU designed the methodology. HA, MK, HA, PX, MT, AH, NOS, and SU conducted a research and investigation process. TI, KM, and SU acquired funds. TI and SU administrated the project. SS and KM supervised the project. HA and SU wrote the original draft, and all the authors participated in writing the final manuscript.

## Competing Interests statement

H.A. reports stock for ImmunoGeneTeqs, Inc. S.S. reports advisory role for ImmunoGeneTeqs, Inc; stock for ImmunoGeneTeqs, Inc, T.I. reports research funding from Rohto Pharmaceutical Co., Ltd., and Biometrics Sympathies Inc. K.M. reports consulting or advisory role for Kyowa-Hakko Kirin, ImmunoGeneTeqs, Inc; research funding from Kyowa-Hakko Kirin, and Ono; stock for ImmunoGeneTeqs, Inc, IDAC Theranostics, Inc. S.U. reports advisory role for ImmunoGeneTeqs, Inc; stock for ImmunoGeneTeqs, Inc, IDAC Theranostics, Inc.

## Notes

### Summary of Updates

We added the data of bulk TCR sequencing up to 6 months after the third vaccination, TCR sequencing on Spike epitope-specific T cells, and single-cell TCR/RNA sequencing on five donors. We also analyzed the temporal kinetics of vaccine-responding clonotypes at an individual epitope level.

