## Supplemental Figures and Tables for "CD8^+^ T-cell memory induced by successive SARS-CoV-2 mRNA vaccinations is characterized by clonal replacement"

### Supplementary Figure S1

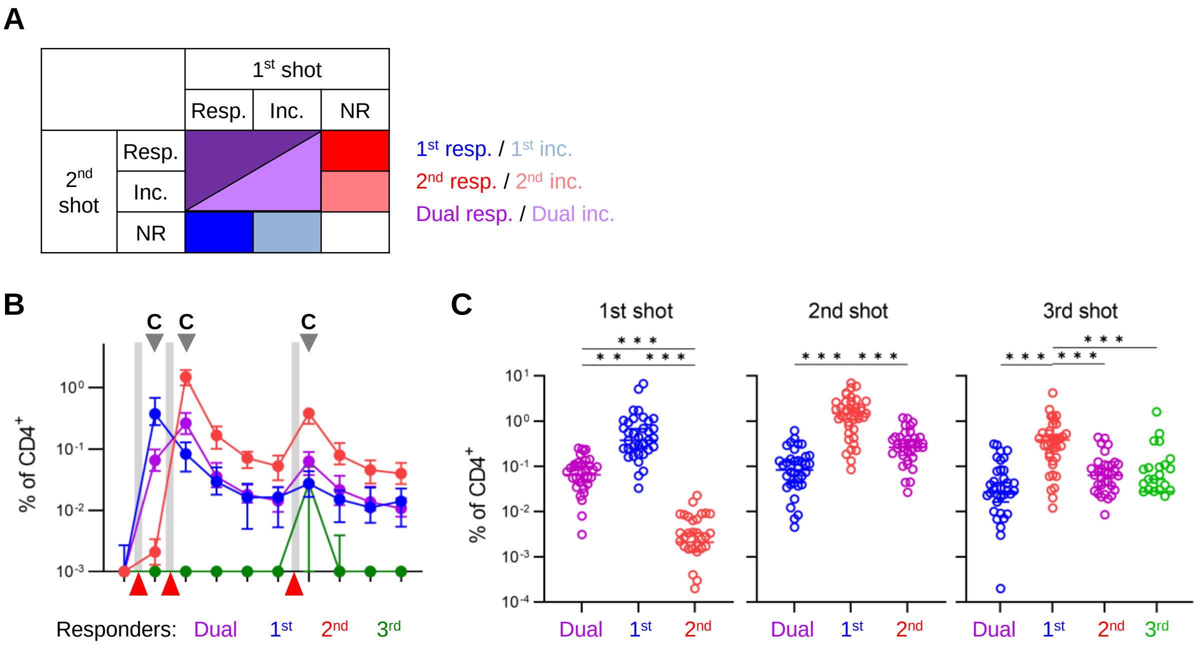


**Fig. S1**. **Kinetics of responding and increasing clones through vaccination.** (A) Overview for identifying 1st, 2nd, and Dual responder clones. Responding (Resp.), increased (Inc.), and non-responsive (NR) clones were identified after each shot of vaccination. 1st, 2nd, and Dual res./inc. clones were defined as indicated. The detail is described in the Materials and Method section. (F) Kinetics of $CD4^+$ 1st, 2nd, Dual, and 3rd responder clone frequencies. (F) The comparison of total frequency of $CD4^+$ 1st, 2nd, Dual, and 3rd responder clonotypes after each shot of vaccination. The total frequency for 3rd responders were plotted only after the third shot (right). (B) Median with 95% CI was plotted. (C) Median was plotted. Friedman test with Dunn's multiple comparisons was performed. n = 38 except for P11 (n=37). ***P<0.001; **P<0.01.

Supplementary Figure S2

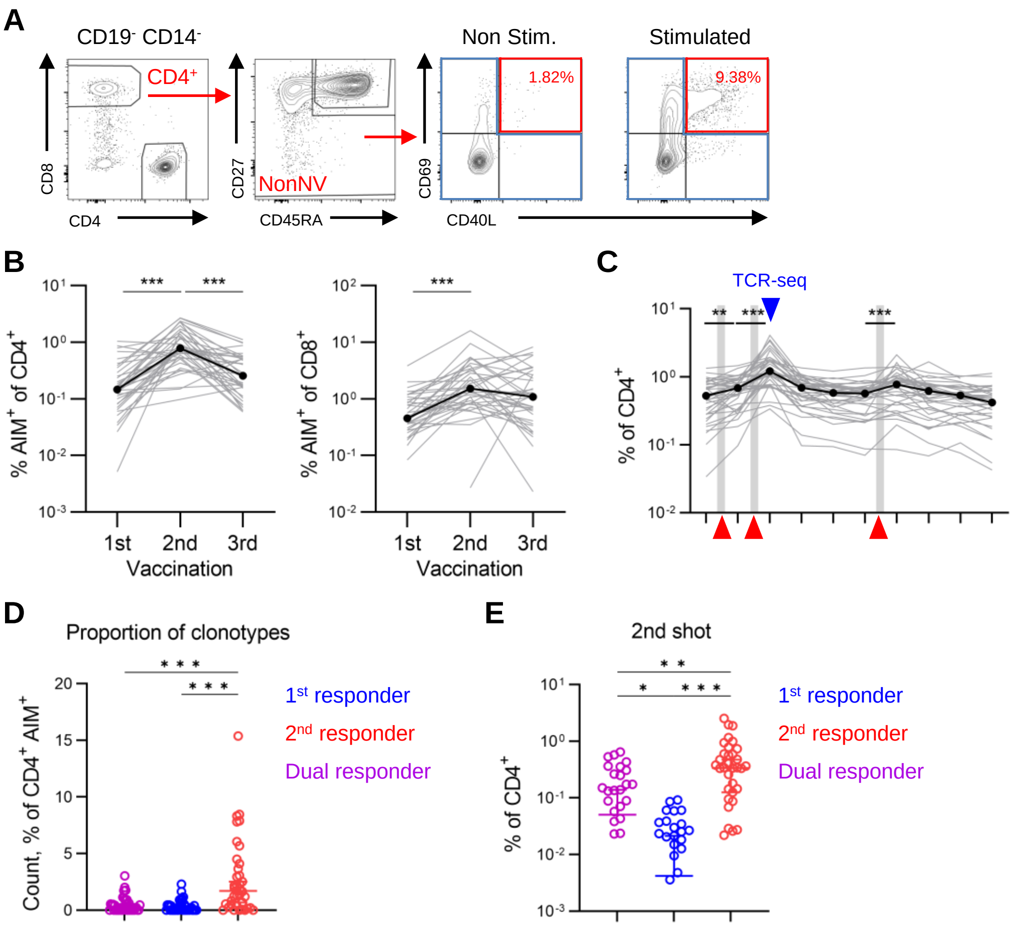


**Fig. S2**. **Response pattern of CD8$^+$ Spike-reactive clonotypes.** (A) Gating strategy for sorting AIM$^+$ and AIM$^-$ non-naive CD4$^+$ T cells. The proportion of AIM$^+$ cells among non-naive CD4$^+$ T cells was also shown. (B) Proportion of AIM+ cells of CD8+ (left) and CD4+ (left) T cells after vaccination. (C) Kinetics of the total frequency of CD4$^+$ AIM$^+$ clonotypes of each participant (thin line, grey). The median was also plotted (thick line, black). (D) The proportion of 1$^{st}$, 2$^{nd}$, and Dual responders in CD4$^+$ AIM$^+$ clonotypes. The count of AIM$^+$ responder clonotypes divided by that of all AIM$^+$ clonotypes was compared. (G) The total frequency of CD4$^+$ AIM$^+$ 1$^{st}$, 2$^{nd}$, and Dual responders after the second shot. (B-E) Friedman test with Dunn's multiple comparisons was performed. n = 38 except for P11 (n=37). ***P<0.001; **P<0.01; *P<0.05.

Supplementary Figure S3

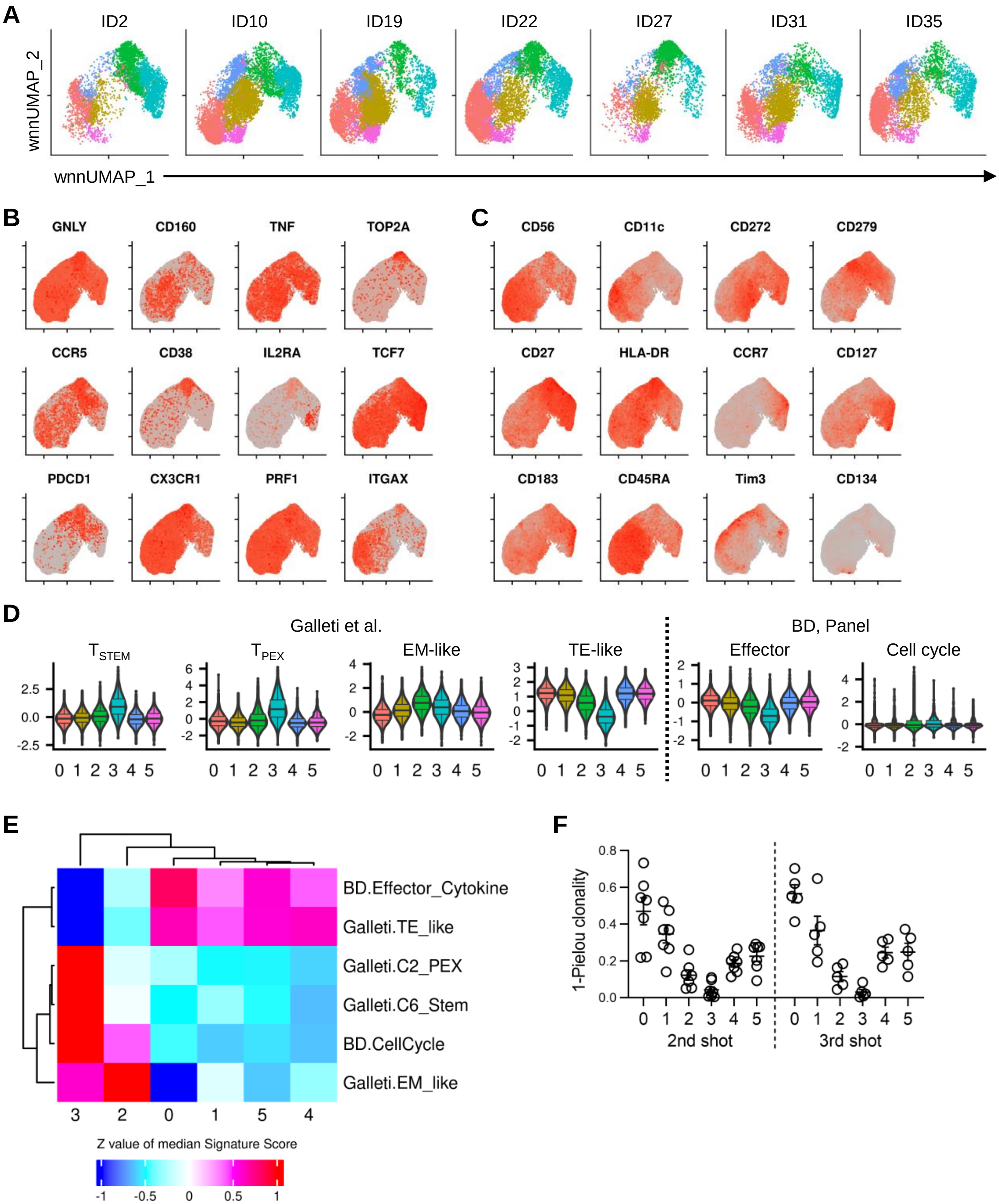


**Fig. S3. Characterization of CD8⁺ T cell clusters observed in single-cell analysis.** (A) wnnUmap plot split by the participants. (B and C) Feature plots showing the mRNA (B) and protein (C) expression of marker genes. (D) Violin plot showing gene signature scores of T-cell clusters. Signature genes are referred from the marker genes of T-cell clusters described in (Galleti) and the datasheet of BD Rhapsody™ Immune Response Panel Hs. (E) Heat map for the Signature Scores of each T-cell clusters. Median of Signature Scores of cells within the cluster was calculated, then normalized between the Signatures. (F) Clonality of the CD8⁺ T-cell clusters.

Supplementary Figure S4

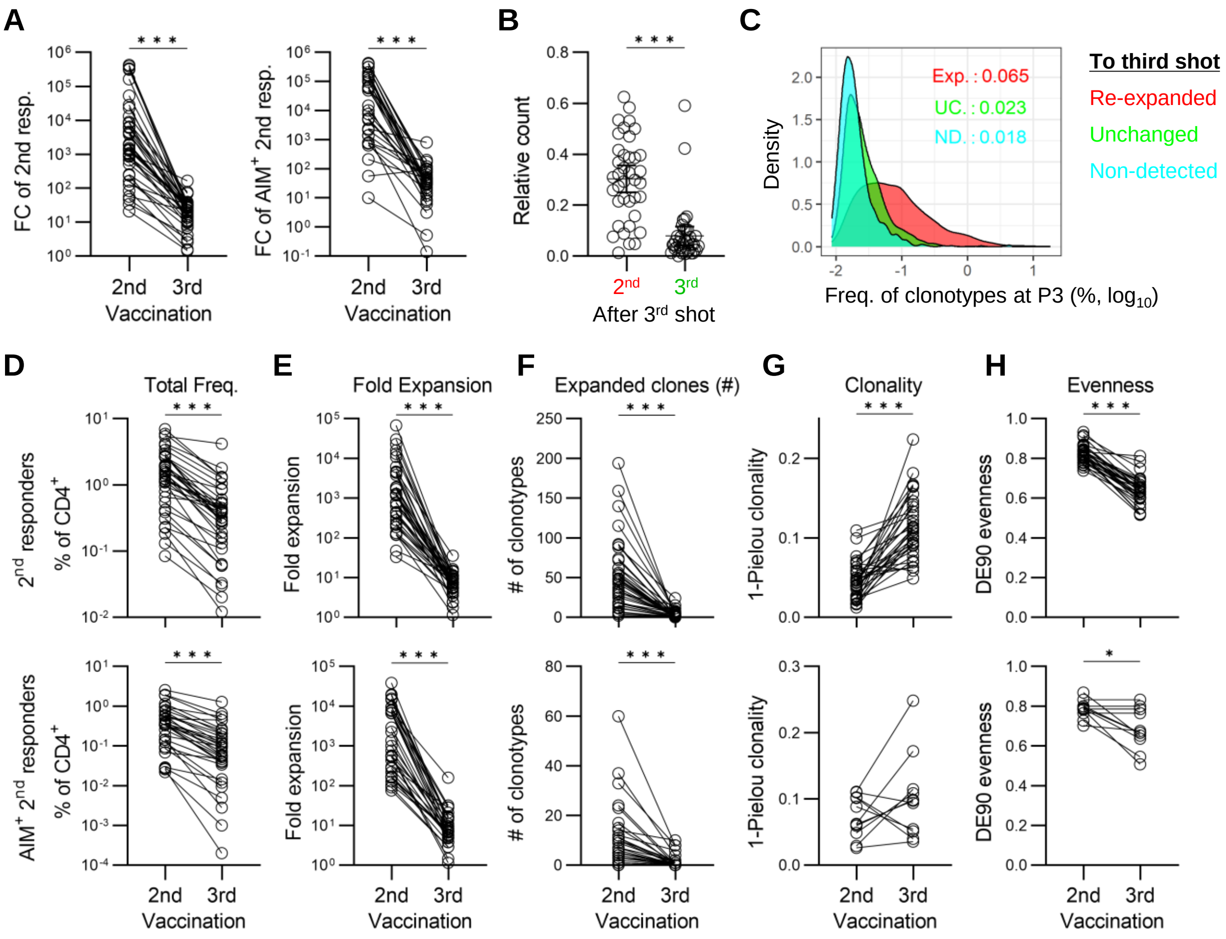


**Fig. S4. Analysis of CD8$^{+}$ and CD4$^{+}$ 2$^{nd}$ responder clonotypes after the third shot.** (A) Fold expansion of CD8$^{+}$ all (left) or AIM$^{+}$ (right) 2$^{nd}$ responder clonotypes after the vaccination. (B) Count of expanding clonotypes after the third shot within CD8$^{+}$ 2$^{nd}$ and 3$^{rd}$ responders relative to that of 2$^{nd}$ responders after the second shot. (C) Density plot of the frequency of 2$^{nd}$ responder clonotypes at P3. 2$^{nd}$ responders were grouped by their response to the third shot: expanded to the third shot (Expanded: red), not expanded but existed after the third shot (Unchanged: green), and not detected after the third shot (Non-detected: blue). (D) Total frequency of CD4$^{+}$ all (top) or AIM$^{+}$ (bottom) 2nd responders after the second or the third shot. (E) Fold expansion of CD4$^{+}$ all (left) or AIM$^{+}$ (right) 2$^{nd}$ responder clonotypes after the vaccination. (F) Count of clonotypes that expanded after the second or the third shot within CD4$^{+}$ all (top) or AIM$^{+}$ (bottom) 2$^{nd}$ responders. (G) Clonality of CD4$^{+}$ all (top) or AIM$^{+}$ (bottom) 2$^{nd}$ responders after the second or the third shot. (H) Evenness of CD4$^{+}$ all (top) or AIM$^{+}$ (bottom) 2$^{nd}$ responders after the second or the third shot. (A, B and D-H) Wilcoxon matched-pairs signed rank test. (A and E) Fold expansion was defined as the total frequency of clonotypes after the vaccination divided by that before the vaccination, which is calculated when at least one responding clonotype presents after the vaccination. (G and H) Clonality and evenness are calculated when more than ten clonotypes exist. n = 38 except for the following; right of (A) (n=32), the bottom of (E) (n=33), and the bottom of (G) and (H) (n=11). ***$P<0.001$.

Supplementary Figure S5

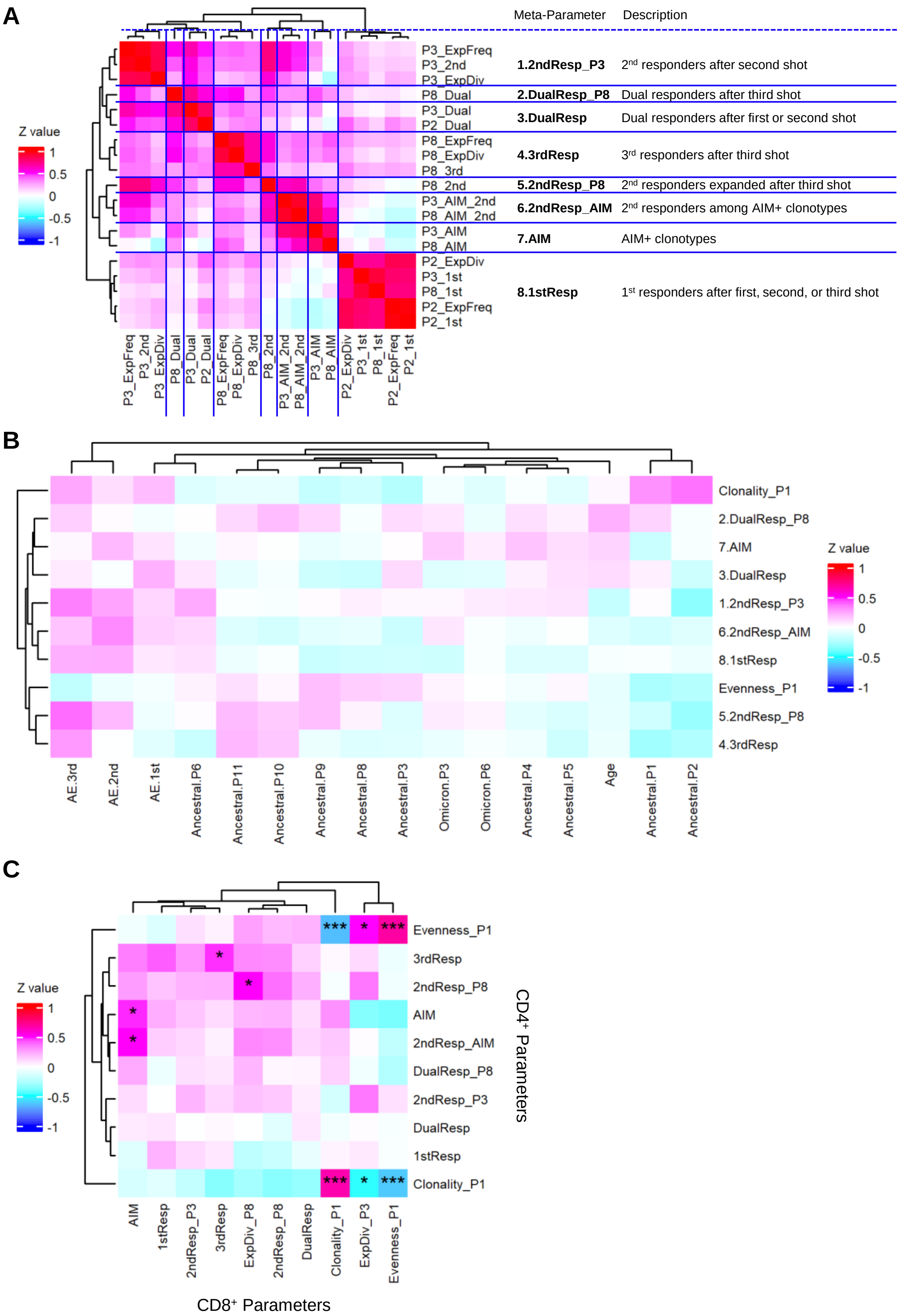

**Fig. S5. Correlation analysis on $CD4^+$ and $CD8^+$ T cell repertoire parameters.** (A) Heatmap of the correlation between $CD4^+$ T cell repertoire parameters. The repertoire meta-parameter is calculated as the average of mutually correlated repertoire parameters (Materials and Methods). The names of repertoire meta-parameters and their descriptions are also shown. (B) Heatmap of the correlation between $CD4^+$ T cell repertoire meta-parameters (row) and the age of participants, the titer of anti-SARS-CoV-2 RBD IgG antibodies ("Ancestral" or "Omicron"), or strength of the adverse events ("AE") (column). © Heatmap of the correlation between $CD4^+$ and $CD8^+$ T cell repertoire meta-parameters. (A-C) Spearman correlation coefficient was colored in heatmap. © q-values of Spearman correlation coefficient between two parameters are plotted. ***q<0.001; *q<0.05.

Supplementary Figure S6

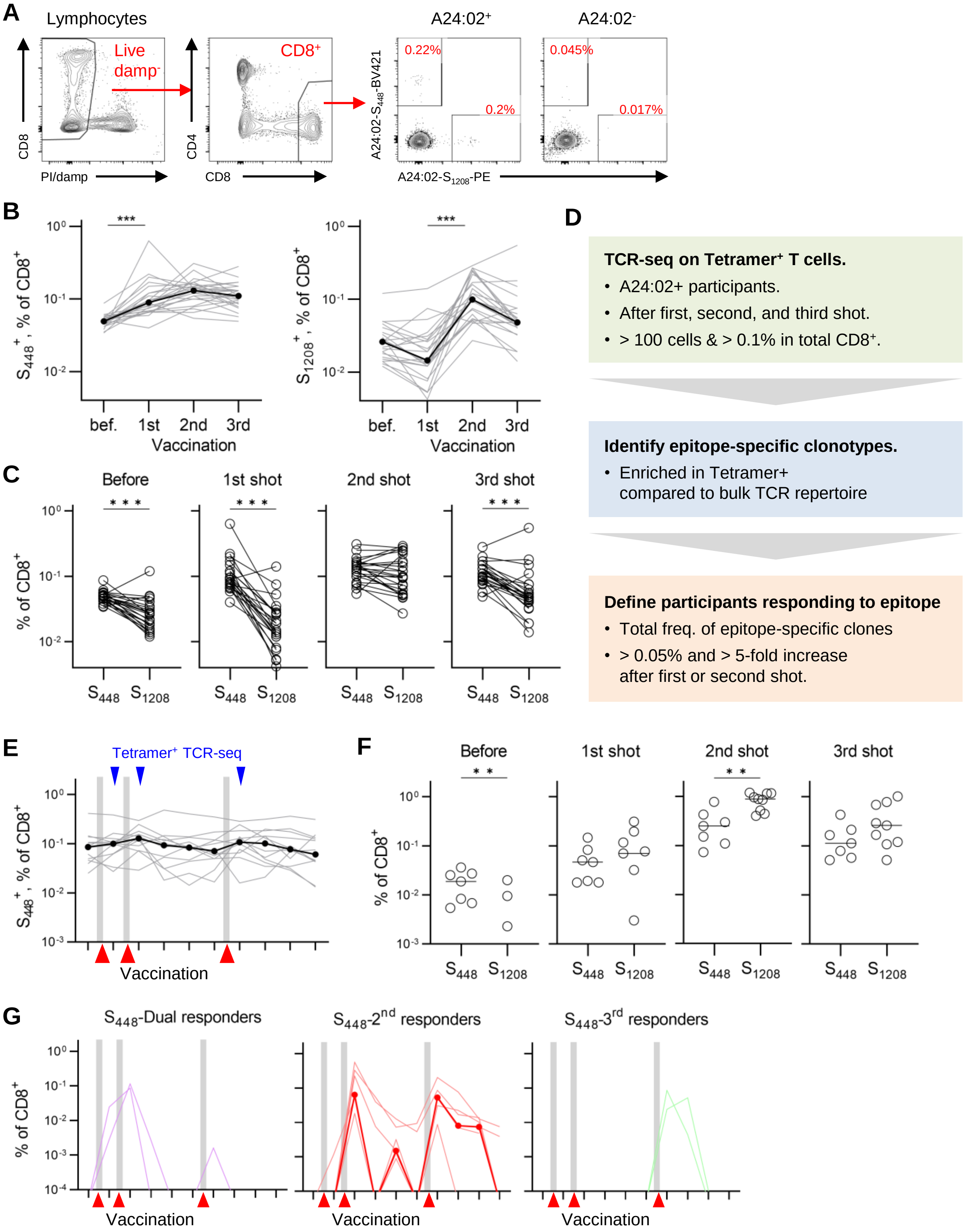

**Fig. S6. Detecting Spike specific clonotypes using pMHC-tetramer staining.** (A) Gating strategy for sorting A24:02-$S_{448}^{+}$ or $S_{1208}^{+}$ $CD8^{+}$ T cells. The proportion of $S_{448}^{+}$ or $S_{1208}^{+}$ cells among non-naive $CD8^{+}$ T cells was also shown. (B) Proportion of $S_{448}^{+}$ (left) or $S_{1208}^{+}$ (right) cells of $CD8^{+}$ T cells after vaccination. (C) Comparison of the proportion of $S_{448}^{+}$ and $S_{1208}^{+}$ cells after vaccination. (D) Flow chart for identifying $S_{448}$ and $S_{1208}$-specific clonotypes. $S_{448}^{+}$ and $S_{1208}^{+}$ cells were sorted using the gating as (A), then TCR-seq was performed when (i) more than 100 cells were sorted and (ii) the proportion of tetramer+ cells was more than 0.1% in $CD8^{+}$ T cells (green). Epitope-specific clonotypes were identified as clonotypes that were enriched in the $tetramer^{+}$ T-cell repertoire (blue). Then, epitope-responsive participants were defined as those whose total frequency of epitope-specific clonotypes was more than 0.05% and 5-fold increased after the first or the second shot (red). (E) Total frequency of $CD8^{+}$ $S_{448}^{+}$ clonotypes of participants who were not epitope-responsive (thin line, grey). The median was also plotted (thick line, black). (F) Comparison of the total frequency of $S_{448}$- and $S_{1208}$-specific clonotypes after vaccination. (G) Kinetics of $S_{448}$-specific 1st, 2nd, Dual, and 3rd responder clonotype frequencies. The thin line represents the total frequency for each participant. The thick line represents the median for $S_{1208}$-responsive participants. The median was calculated only when responder clonotypes were detected in more than half of the participants. (B and C) n = 24. ID22 with high background in $S_{1208}$-$tetramer^{+}$ sample was excluded from the analysis. (E) n = 12. (F) n = 7 or 9 for $S_{448}$- or $S_{1208}$- responsive participants, respectively. (B and E) Friedman test with Dunn's multiple comparisons was performed. (C) Wilcoxon matched-pairs signed rank test was performed. (F) Mann-Whitney test was performed. ***P<0.001; **P<0.01; *P<0.05.

### Supplementary Figure S7

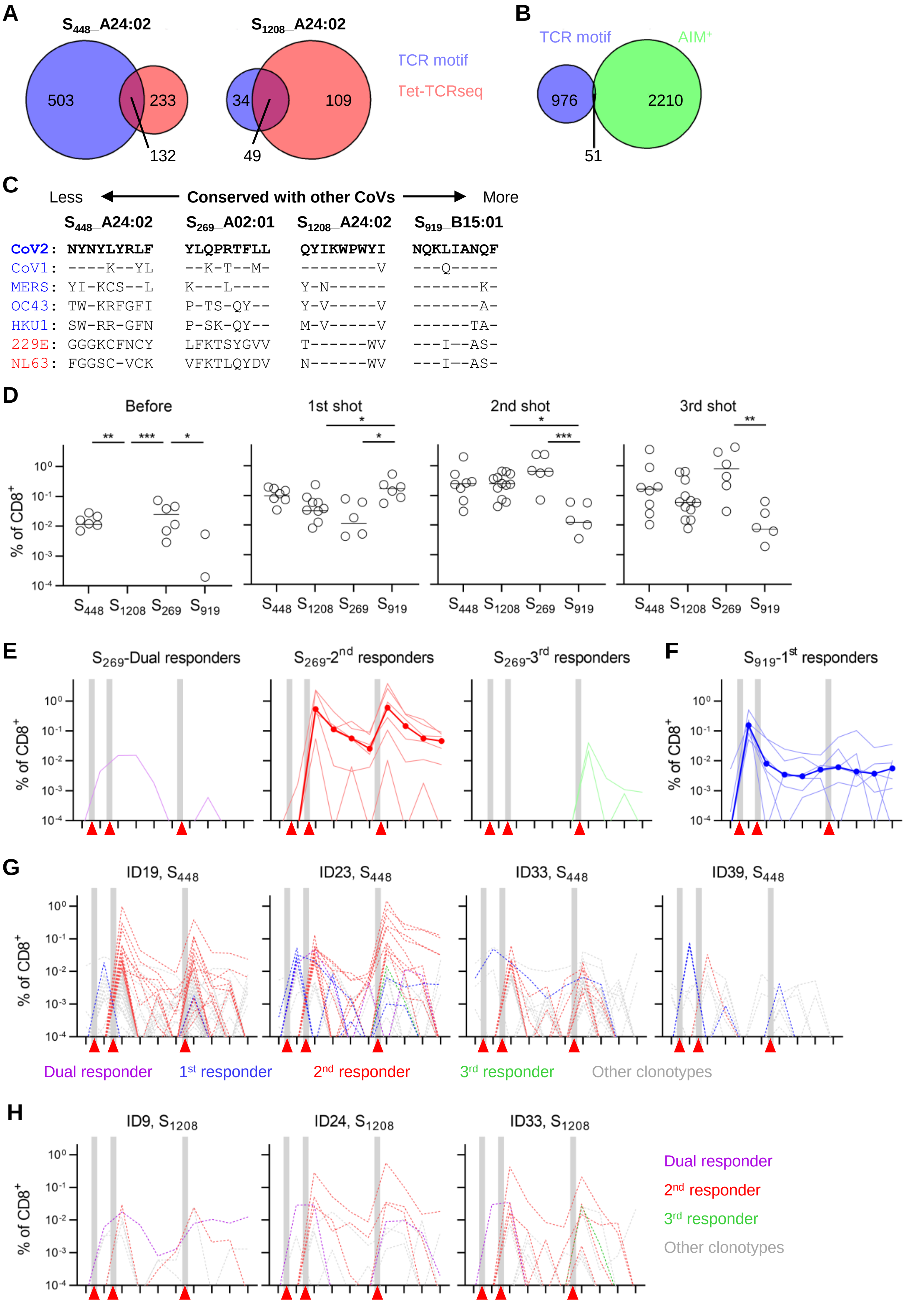

**Fig. S7 Detecting Spike-specific clonotypes using TCR sequence motifs.** (A) Amino acid sequence of Spike epitopes for which TCR sequence motif was detected in our dataset. Their homology with other human coronaviruses is also shown. (B) Venn diagram showing overlap of $S_{448}$- (left) or $S_{1208}$- (right) specific clonotypes detected by TCR motif analysis (blue) and tetramer-TCRseq (red). (C) Venn diagram showing overlap of Spike-specific clonotypes detected by TCR motif analysis ($S_{448}$, $S_{1208}$, $S_{269}$, and $S_{919}$: blue) and AIM$^{+}$ clonotypes (green). (D) Comparison of the total frequency of clonotypes specific to each Spike epitope before and after vaccination. (E and F) Kinetics of $S_{269}$-specific $2^{nd}$, Dual, and $3^{rd}$ responder (E) or $S_{919}$-specific $1^{st}$ responder (F) clonotype frequencies. The thin line represents the total frequency for each participant. The thick line represents the median for epitope-responsive participants. (G and H) Longitudinal tracking of individual $S_{1208}$- (H) and $S_{448}$- (I) specific clonotypes (dashed lines) in some participants. Clonotypes were plotted by response patterns ($1^{st}$ responders: blue, $2^{nd}$ responders: red, Dual responders: purple, $3^{rd}$ responders: green, and other clonotypes: gray). (D) Kruskal-Wallis test with Dunn's multiple comparisons was performed. ***$P<0.001$; **$P<0.01$; *$P<0.05$.

Supplementary Table S2. Metadata of participants on whom single-cell analysis was performed

| ID | Age | Gender | Infection | 2nd shot※ | | 3rd shot※ |
|---|---|---|---|---|---|---|
| 2 | 43 | M | Pre | CD4 | CD8 | |
| 10 | 41 | M | not | | CD8 (ST1※※) | CD8 (ST6) |
| 19 | 35 | F | not | | CD8 (ST2) | CD8 (ST7) |
| 22 | 34 | M | not | | CD8 (ST3) | CD8 (ST8) |
| 27 | 35 | M | not | CD4 | CD8 | |
| 31 | 36 | M | not | | CD8 (ST4) | CD8 (ST9) |
| 35 | 36 | F | not | | CD8 (ST5) | CD8 (ST10) |

※ Colors represent the batch of single-cell analysis.

※※ Name of Sample Tag used for sample preparation. Sample Tags were not used for samples from ID2 and ID27 because these samples were not multiplexed.

Supplementary Table S3. Reagent list used in this study

| Name of reagent | Source | Identifier |
|---|---|---|
| BD vacutainer CPT tubes | BD | #362753 |
| CELLBANKER 1 | Takara | #CB011 |
| RayBio COVID-19 S1 RBD protein Human IgG ELISA Kit | RayBiotech | #IEQ-CoVS1RBD-IgG |
| RayBio COVID-19 Omicron Spike Variant Human IgG ELISA Kit | RayBiotech | IEQ-CoVSM2-IgG |
| RayBio COVID-19 N protein Human IgG ELISA Kit | RayBiotech | IEQ-CoVN-IgG |
| Albumin, Bovine Serum, General Grade, pH7.0 | Nacarai | #01860-65 |
| 0.5M EDTA pH8.0 | NIPPON GENE | #311-90075 |
| D-PBS(-) | Nacarai | #14249-24 |
| BD IMag™ Anti-PE Magnetic Particles (Clone E31-1459) | BD | #557899 |
| DynaMag-96 SideSkirted | Veritas | #DB12027 |
| BD IMag Streptavidin Particles Plus-DM | BD | #557812 |
| Flow-Count Fluorospheres | Beckman | #7547053 |
| Geno Plus Genomic DNA Extraction Miniprep System | VIOGENE | #GG2002 |
| FBS(Netherland) | MP Bio Japan | #2917354H<br>(Lot# #N0218) |
| 1mol/l-HEPES Buffer Solution | Nacarai | #17557-94 |
| Penicillin-Streptomycin Mixed Solution(Stabilized) | Nacarai | #09367-34 |
| BD FastImmune™ Co-Stimulatory Antibodies (CD28/CD49d) | BD | #347690 |
| PepTivator SARS-CoV-2 Prot_S Complete, research grade | Miltenyi | #130-127-951 |
| Propidium iodide | Sigma-Aldrich | #537059 |
| Cell Staining Buffer | BioLegend | #420201 |
| BD® AbSeq Immune Discovery Panel | BD | #625970 |
| Ms Single Cell Sample Multiplexing Kit | BD | #633793 |
| $S_{448}$ peptide (NYNYLYRLF, purity = 99.2%) | GenScript | |
| $S_{1208}$ peptide (QYIKWPWYI, purity = 91.1%) | GenScript | |
| QuickSwitch™ Quant HLA-A*24:02 Tetramer Kit-BV421 | MBL | #TB-7302-K4 |
| QuickSwitchTM Quant HLA-A*24:02 Tetramer Kit-PE | MBL | #TB-7302-K1 |
| Dasatinib | AdooQ | A10290-25 |
| Dynabeads M-270 Streptavidin | Invitrogen | #DB65306 |
| Deoxy-NTP Set | Roche | #11969064001 |
| Betaine solution 5 M, PCR Reagent | Sigma-Aldrich | #B0300-5VL |
| 1M MgCl2 | NIPPON GENE | #310-90361 |
| RNasin Plus Ribonuclease Inhibitor | Promega | #N2611 |
| SuperScript II Reverse Transcriptase | Invitrogen | #18064071 |
| KAPA HiFi HS ReadyMix | KAPA Biosystems | #KK2602 |
| AMPure XP | Beckman | #A63881 |
| NovaSeq 6000 S4 Reagent Kit v1.5 | Illumiina | #20028313 |
| KAPA SYBR Fast qPCR Kit | KAPA Biosystems | #KK4602 |
| Calcein AM | Nacarai | #06735-81 |
| BD Rhapsody Cartridge Kit | BD | #633733 |
| BD Rhapsody™ Cartridge Reagent Kit | BD | #633731 |
| BD Rhapsody cDNA Kit | BD | #633773 |
| Targeted mRNA and AbSeq Amplification Kit | BD | #633774 |
| Immune Response Panel HS | BD | #633750 |
| NEBuffer 2 | NEB | #B7002 |
| Klenow Fragment (3'→5' exo-) | NEB | #M0212M |
| ProNex Size-Selective Purification System | Promega | #NG2001 |

Supplementary Table S4. Antibody list used in this study

| Name of antibody | Source | RRID or Catalog No. | Antibody Mix | | | |
|---|---|---|---|---|---|---|
| | | | 1 | 2 | 3 | 4 |
| Human TruStain FcX™ (Fc Receptor Blocking Solution) | BioLegend | AB_2818986 | O | | | |
| Mouse anti-human CD3 Antibody (Pacific Blue) (clone SK7) | BioLegend | AB_2563422 | O | | O | |
| Mouse anti-human CD4 Antibody (PE) (clone SK3) | BioLegend | AB_1937246 | O | | O | |
| Mouse anti-human CD8 Antibody (APC) (clone SK1) | BioLegend | AB_2075388 | O | | | |
| Mouse anti-human CD19 Antibody (APC) (clone HIB19) | BD | AB_398597 | O | | | |
| Mouse anti-APC Antibody (Biotin) (clone APC003) | BioLegend | AB_345360 | | | | |
| Mouse anti-human CD40 Antibody (Purified) (clone HB14) | BioLegend | AB_314965 | | | | |
| Mouse anti-human CD107a (LAMP-1) Antibody (FITC) (clone H4A3) | BioLegend | AB_1186036 | | | | |
| Mouse anti-human CD4 Antibody (PerCP/Cyanine5.5) (clone SK3) | BioLegend | AB_1953236 | | O | | O |
| Mouse anti-human CD8 Antibody (Brilliant Violet 510) (clone RPA-T8) | BioLegend | AB_2561942 | | O | | |
| Mouse anti-human CD14 Antibody (PE/Dazzle™ 594) (clone 63D3) | BioLegend | AB_2750398 | | O | O | O |
| Mouse anti-human CD19 Antibody (PE/Dazzle™ 594) (clone HIB19) | BioLegend | AB_2563560 | | O | O | O |
| Mouse anti-human CD45RA Antibody (APC/Cyanine7) (clone HI100) | BioLegend | AB_10708880 | | O | O | O |
| Mouse anti-human CD27 Antibody (PE/Cyanine7) (clone O323) | BioLegend | AB_2561918 | | O | O | |
| Mouse anti-human CD154 Antibody (APC) (clone 24-31) | BioLegend | AB_314832 | | O | | |
| Mouse anti-human CD69 Antibody (Pacific Blue) (clone FN50) | BioLegend | AB_493666 | | O | | |
| Mouse anti-human CD137 Antibody (PE) (clone 4B4-1) | BD | AB_10926375 | | O | | |
| Mouse anti-human CD8 Antibody (FITC) (clone SK1) | BioLegend | AB_1877178 | | | O | |
| Mouse anti-human CD8 Antibody (FITC) (clone HIT8a) | BioLegend | AB_314110 | | | | O |
| Mouse anti-human CD27 Antibody (APC) (clone O323) | BioLegend | AB_314302 | | | | O |

Supplementary Table S5. Software and algorithms used in this study

| **Name of software and algorithms** | **Source** | **Identifier** |
|---|---|---|
| FlowJo (v10.8.1) | BD<br>RRID:SCR_008520 | https://www.flowjo.com/ |
| BD FACSDiva Software (v8.0.1) | BD<br>RRID:SCR_001456 | http://www.bdbiosciences.com/instruments/software/facsdiva/index.jsp |
| Cutadapt (v2.10, 3.2, and 3.4) | (Martin, 2011)<br>RRID:SCR_011841 | https://cutadapt.readthedocs.io/en/v3.2/ |
| PRINSEQ (v0.20.4) | (Schmieder, 2011)<br>RRID:SCR_005454 | https://apolo-docs.readthedocs.io/en/latest/software/applications/prinseq-lite/prinseq-lite-0.20.4/index.html |
| MiXCR (v3.0.5) | (Bolotin, 2015)<br>RRID:SCR_018725 | https://mixcr.readthedocs.io/en/develop/ |
| VDJtools (v1.2.1) | (Shugay, 2015) | https://vdjtools-doc.readthedocs.io/en/master/ |
| bowtie2 (v2.4.2) | (Langmead, 2012)<br>RRID:SCR_016368 | http://bowtie-bio.sourceforge.net/bowtie2/index.shtml |
| pysam (v0.15.4) | (Li, 2009)<br>RRID:SCR_021017 | https://github.com/pysam-developers/pysam |
| DropletUtils (v1.14.1) | (Lun, 2018) | https://bioconductor.org/packages/release/bioc/html/DropletUtils.html |
| Seqkit (v0.15.0) | (Shen, 2016)<br>RRID:SCR_018926 | https://github.com/shenwei356/seqkit/releases |
| Seurat (v4.0.1) | (Hao, 2021)<br>RRID:SCR_007322 | https://satijalab.org/seurat/ |
| ADTnorm | (Zheng, 2022) | https://github.com/yezhengSTAT/ADTnorm |
| lighter (v1.1.2) | (Song, 2014) | https://github.com/mourisl/Lighter |
| R (v4.2.1) | | https://www.r-project.org/ |
| Rstudio (v2022.12.0) | RRID: SCR_000432 | https://posit.co/download/rstudio-desktop/ |
| Docker (v20.10.21) | RRID: SCR_016445 | https://www.docker.com/products/docker-desktop/ |
| GraphPad Prism (v9.5.1) | GraphPad<br>RRID:SCR_002798 | https://www.graphpad.com/scientific-software/prism/ |

Supplementary Table S6. Oligonucleotides list used in this study

| Name of oligonucleotides | Source |
|---|---|
| Bulk TCRseq primer: BioEcoP-dT25-adapter: [BiotinTEG]-AAGCAGTGGTATCAACGCAGAGTACTTTTTTTTTTTTTTTTTTTTTTTTT | (Aoki et al., 2019) |
| Bulk TCRseq primer: i5-TSO: /5AmMC12/ACACGACGCTCTTCCGATCTrGrG+G | (Tsunoda et al., 2021) |
| Bulk TCRseq primer: illumina-i5: ACACGACGCTCTTCCGATCT | (Tsunoda et al., 2021) |
| Bulk TCRseq primer: hTrac_ex: GGAATAATGCTGTTGTTGAAGGCGTTTGC | This study |
| Bulk TCRseq primer: hTrbc_ex: GTGCACCTCCTTCCCATTCACCC | This study |
| Bulk TCRseq primer: illumina-i5_2nd: CACTCTTTCCCTACACGACGCTCTTCCGATCT | This study |
| Bulk TCRseq primer: illumina-i7-[BCX]-hTrbc: GTGACTGGAGTTCAGACGTGTGCTCTTCCGATCT XXXXXXXXXX GGTGGGAACACSTTKTTCAGGTCCT | This study |
| Bulk TCRseq primer: i5_UDI: AATGATACGGCGACCACCGAGATCTACAC XXXXXXXX CACTCTTTCCCTACACGAC | (Shichino et al., 2022) |
| Bulk TCRseq primer: i7_UDI: CAAGCAGAAGACGGCATACGAGAT XXXXXXXX GTGACTGGAGTTCAGACGTGT | (Shichino et al., 2022) |
| Single-cell Sequencing primer: NH2-dT25-TSO: NH2-C6- TTTTTTTTTTTTTTTTTTTTTTTTTrGrGrG | (Aoki et al., 2023) |
| Single-cell Sequencing primer: Universal Oligo-long: NH2-ACACTCTTTCCCTACACGACGCTCTTCCGATCT | (Shichino et al., 2022) |
| Single-cell Sequencing primer: BD_hTRAC_primer1: CTGGAATAATGCTGTTGTTGAAGG | This study |
| Single-cell Sequencing primer: BD_hTRBC_primer1: AGCCCGTAGAACTGGACTT | This study |
| Single-cell Sequencing primer: BD_hTRAC_primer2: ATCAAAATCGGTGAATAGGCAGAC | This study |
| Single-cell Sequencing primer: BD_hTRBC_primer2: GATCTCTGCTTCTGATGGCTCA | This study |
| Single-cell Sequencing primer: illumina-i7-hTrac: GTGACTGGAGTTCAGACGTGTGCTCTTCCGATCTACGGCAGGGTCAGGGTTCTGGATA | This study |
| Single-cell Sequencing primer: illumina-i7-hTrbc: GTGACTGGAGTTCAGACGTGTGCTCTTCCGATCTGGTGGGAACACSTTKTTCAGGTCCT | This study |

Bio: Biotin. TEG: Triethylene glycol. 5AmMC12: 5' Amino Modifier C12. BC: Barcode. rG: Ribonucleic guanine. +G: Affinity Plus modified guanine. UDI: Unique dual index.

Table S7. Reagents and conditions for bulk TCR-seq library preparation

| | Reagents (Table S2) | Primers (Table S5) | Thermal cycler program | | |
|---|---|---|---|---|---|
| RNA trap | Dynabeads M270 streptavidin | BioEcoP-dT25-adapter | | | |
| Reverse Transcription | Superscript II<br>First Strand buffer[※1]<br>DTT[※1]<br>dNTP<br>$MgCl_2$<br>Betain<br>RNaseIn Plus RNase Inhibitor | i5-TSO | 42 ºC | 60 min | |
| First TCR-PCR | KAPA Hifi Hotstart ReadyMix | illumina-i5<br>hTrac_ex<br>hTrbc_ex | 95 ºC<br>98 ºC<br>65 ºC<br>72 ºC<br>72 ºC | 3 min<br>20 sec<br>15 sec<br>30 sec<br>2 min | 10 cycles (98 ºC – 72 ºC 30 sec) |
| Second TCR-PCR | KAPA Hifi Hotstart ReadyMix | illumina-i5_2nd<br>i7-BC[X]_hTrbc | 95 ºC<br>98 ºC<br>65 ºC<br>72 ºC<br>72 ºC | 3 min<br>20 sec<br>15 sec<br>30 sec<br>2 min | 10 cycles [※2, ※3] (98 ºC – 72 ºC 30 sec) |
| Third TCR-PCR | KAPA Hifi Hotstart ReadyMix | i5_UDI<br>i7_UDI | 95 ºC<br>98 ºC<br>65 ºC<br>72 ºC<br>72 ºC | 3 min<br>20 sec<br>15 sec<br>30 sec<br>2 min | 15 cycles [※2, ※3] (98 ºC – 72 ºC 30 sec) |

※1 Included in the SuperScript II kit.
※2 Sample of $AIM^+$ non-naïve $CD4^+$ or $CD8^+$ T cells; 2nd PCR: 10 cycles, 3rd PCR: 18 cycles
※3 Sample of $Tetramer^+$ $CD8^+$ T cells; 2nd PCR: 15 cycles, 3rd PCR: 17 cycles

Table S8. Conditions for PCR reaction and purification in scTCR library preparation

| Reaction | Target | Primers (Table S5) | | Pronex selection[A] | |
|---|---|---|---|---|---|
| | | **Forward** | **Reverse** | **Upper** | **Lower** |
| 1st PCR | Sample Tag | Sample Tag PCR1 primer | Universal Oligo-long | 1:1.25 | 1:1.75 |
| | AbSeq | BD™ AbSeq Primer | Universal Oligo-long | 1:1.25 | 1:1.75 |
| | mRNA | BD immune response panel Hs PCR1 | Universal Oligo-long | | 1:1.25 |
| | TCR | hTRAC_primer1<br>hTRBC_primer1 | Universal Oligo-long | | 1:1.25 |
| 2nd PCR | Sample Tag | SampleTag PCR2 primer | Universal Oligo-long | 1:1.24 | 1:1.5 |
| | mRNA | BD immune response panel Hs PCR2 | Universal Oligo-long | 1:1.05 | 1:1.3 |
| | TCR | BD_hTRAC_primer2<br>BD_hTRBC_primer2 | Universal Oligo-long | 1:0.9 | 1:1.25 |
| 3rd PCR | Abseq | illumina i7-long | Universal Oligo-long | 1:1.24 | 1:1.5 |
| | mRNA | illumina i7-long | Universal Oligo-long | 1:1.05 | 1:1.3 |
| | TCR | illumina-i7-hTrac<br>illumina-i7-hTrbc | Universal Oligo-long | 1:0.9 | 1:1.25 |
| 4tf PCR | Sample Tag | i5_UDI | i7_UDI | 1:1.24 | 1:1.5 |
| | Abseq | i5_UDI | i7_UDI | 1:1.24 | 1:1.5 |
| | mRNA | i5_UDI | i7_UDI | 1:1.05 | 1:1.3 |
| | TCR | i5_UDI | i7_UDI | 1:0.9 | 1:1.25 |

[A] Sample-to-beads ratios for purifying PCR product using Pronex beads.

Table S9. Parameters for MiXCR to analyze bulk TCRseq data

| Name of parameter | Settings |
|---|---|
| --starting-material | rna |
| --5-end | no-v-primers |
| --3-end | c-primers |
| --adapters | no-adapters |
| --receptor-type | Trb |
| --region-of-interest | CDR3 |
| --only-productive | |
| --align | -OvjAlignmentOrder=JThenV |
| --align | -OvParameters.geneFeatureToAlign=VTranscript |
| --assemble | -ObadQualityThreshold=15 |
| --assemble | -OseparateByV=true |
| --assemble | -OseparateByJ=true |

Table S10. Genes used for calculating signature scores

| Signature | Genes |
|---|---|
| $T_{STEM}$ | *PASK, LEF1, LTB, SELL, TNFSF10, DUSP4, CD52, FYB, FTH1, LGALS1, JUNB, BIRC3, S100A10, CCR7, TRAT1, CD8B, IL4R, FOXP1, RGS1* |
| $T_{PEX}$ | *CCR7, CD8B, LEF1, PASK, FYB, SELL, FOXP1, TCF7, KLRC1* |
| EM-like | *RGS1, CD74, GZMK, TIGIT, CD8B, CCL5, CST7, HLA-DPA1, PTPRC, CCL4, HLA-DRA, JUN, JUNB, HLA-DQB1, HLA-DMA, CD27, DUSP2, IFNG, FYN* |
| TE-like | *GZMH, GZMB, GNLY, FCGR3A, NKG7, CCL4, CST7, ITGB2, LAIR2, CTSW, CCL5, CX3CR1, KLRF1, GZMA, CD3D, PRF1, LGALS1, HLA-DPA1, CD52, CD8A, CD3G, APOBEC3G, PTPRC, IFNG, CD8B, IFITM2, TIGIT, CD300A, CD63, CD160, CD2, CTSD, BIN2* |
| Effector | *CMTM2, CSF2, CSF3, FASLG, GNLY, GZMA, GZMB, GZMH, GZMK, IFNA1, IFNG, LIF, LTA, LTB, NAMPT, PRF1, SPP1, TNF, TNFSF10, TNFSF13, TNFSF13B, TNFSF14, TNFSF8* |
| Cell cycle | *AURKB, CCND2, HMMR, KIAA0101, MCM2, MCM4, MKI67, MYC, PCNA, TOP2A, TXK, TYMS, UBE2C* |

Supplementary Table S11. Parameters for MiXCR to analyze single-cell TCRseq data

| Name of parameter | Settings |
|---|---|
| --starting-material | rna |
| --5-end | no-v-primers |
| --3-end | c-primers |
| --adapters | no-adapters |
| --receptor-type | tra or trb |
| --region-of-interest | CDR3 |
| --only-productive | |
| --contig-assembly | |
| --align | -OvjAlignmentOrder=JThenV |
| --align | -OvParameters.geneFeatureToAlign=VTranscriptWithout5UTRWithP |
| --align | -OvParameters.parameters.mapperKValue=5 |
| --align | -OjParameters.parameters.mapperKValue=5 |
| --align | -OcParameters.parameters.mapperKValue=5 |
| --align | -OvParameters.parameters.floatingLeftBound=true |
| --align | -OjParameters.parameters.floatingRightBound=false |
| --align | -OvParameters.parameters.maxAdjacentIndels=2 |
| --align | -OjParameters.parameters.maxAdjacentIndels=2 |
| --align | -OcParameters.parameters.maxAdjacentIndels=2 |
| --align | -OvParameters.parameters.absoluteMinScore=40 |
| --align | -OjParameters.parameters.absoluteMinScore=40 |
| --align | -OcParameters.parameters.absoluteMinScore=15 |
| --align | -OvParameters.parameters.relativeMinScore=0.87 |
| --align | -OjParameters.parameters.relativeMinScore=0.87 |
| --align | -OcParameters.parameters.relativeMinScore=0.87 |
| --align | -OdParameters.absoluteMinScore=20 |
| --align | -OdParameters.relativeMinScore=0.7 |
| --align | -OallowPartialAlignments=true |
| --align | -OminSumScore=120 |
| --assemble | -ObadQualityThreshold=15¥ |
| --assemble | -OseparateByV=false |
| --assemble | -OaddReadsCountOnClustering=true |
| --assemble | -OseparateByC=false |
| --assemble | -OseparateByJ=false |
| --assemble | -OcloneClusteringParameters.searchDepth=50 |
| --assemble | -OcloneClusteringParameters.allowedMutationsInNRegions=4 |
| --assemble | -OcloneClusteringParameters.searchParameters=fourMismatchesOrIndels |
| --assemble | -OcloneClusteringParameters.clusteringFilter.specificMutationProbability=1E-2 |
| --assembleContigs | -OsubCloningRegion=CDR3 |
| --assembleContigs | -OalignedRegionsOnly=false |
| --export | -p fullImputed |
| --export | -vHits |
| --export | -vHit |
| --export | -dHits |
| --export | -dHit |
| --export | -jHits |
| --export | -jHit |
| --export | -nFeatureImputed L+FR2+CDR2+FR3+CDR3+FR4+CDR4+FR4 |
| --export | -aaFeatureImputed L+FR2+CDR2+FR3+CDR3+FR4+CDR4+FR4 |
| --export | --minimal-clone-fraction 0.01 |
